# SpotClean adjusts for spot swapping in spatial transcriptomics data

**DOI:** 10.1101/2021.06.11.448105

**Authors:** Zijian Ni, Aman Prasad, Shuyang Chen, Richard B. Halberg, Lisa Arkin, Beth Drolet, Michael Newton, Christina Kendziorski

## Abstract

Spatial transcriptomics (ST) is a powerful and widely-used approach for profiling genome-wide gene expression across a tissue with emerging applications in molecular medicine and tumor diagnostics. Recent spatial transcriptomics experiments utilize slides containing thousands of spots with spot-specific barcodes that bind mRNA. Ideally, unique molecular identifiers at a spot measure spot-specific expression, but this is often not the case owing to bleed from nearby spots, an artifact we refer to as spot swapping. We propose SpotClean to adjust for spot swapping and, in doing so, to increase the sensitivity and precision with which downstream analyses are conducted.

---

Spatial transcriptomics (ST) is a powerful and widely-used approach for profiling genome-wide gene expression across a tissue ^1,2^. In a typical ST experiment, fresh-frozen (or FFPE) tissue is sectioned and placed onto a slide containing spots, with each spot containing millions of capture oligonucleotides with spatial barcodes unique to that spot. The tissue is imaged, typically via Hematoxylin and Eosin (H&E) staining. Following imaging, the tissue is permeabilized to release mRNA which then binds to the capture oligonucleotides, generating a cDNA library consisting of transcripts bound by barcodes that preserve spatial information. Data from an ST experiment consists of the tissue image coupled with RNA-sequencing data collected from each spot. A first step in processing ST data is tissue detection, where spots on the slide containing tissue are distinguished from background spots without tissue. Unique molecular identifier (UMI) counts at each spot containing tissue are then used in downstream analyses (Supplementary Figure 1).

**Figure 1:**
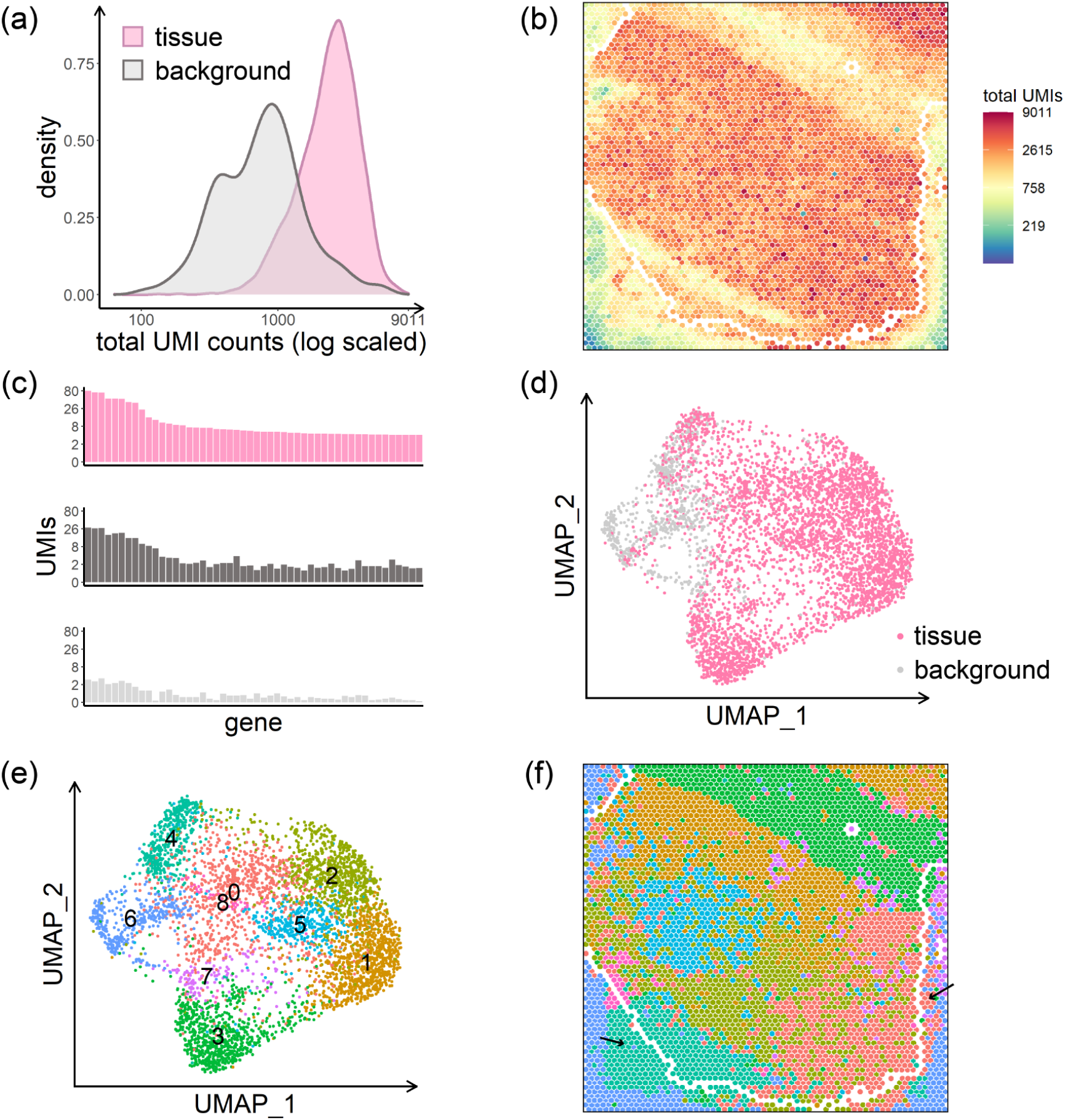
Data from the human dorsolateral prefrontal cortex profiled in the spatialLIBD experiment, sample LIBD_151507. (a) UMI count densities for tissue and background spots show relatively high counts in the background. (b) UMI total counts in the background decrease with increasing distance from the tissue; the perimeter delineating tissue and background is shown in white. (c) Counts of the top 50 genes from a select tissue region (upper), from a nearby background region (middle), and from a distant background region (bottom) show the similarity between expression in tissue spots and nearby background spots due to spot swapping from tissue to background, an effect that decreases as distance from the tissue increases. The positions of the three regions are shown in Supplementary Figure 2. (d) Tissue and background spots are not distinguished visually via UMAP. (e) Graph-based clustering of all spots identifies 9 clusters. (f) Spots on the slide are colored by their cluster membership shown in (e). Black arrows highlight areas of spot swapping of signal from tissue to background. Spots on the perimeter (shown in white) have been removed from the summaries shown here to ensure that the effects shown are not due to spots on the tissue-background boundary. The H&E image for this dataset is shown in Supplementary Figure 2.

Ideally, a gene-specific UMI at a given spot would represent expression of that gene at that spot, and spots without tissue would show no UMIs. This is not the case in practice. Messenger RNA bleed from nearby spots causes substantial contamination of UMI counts, an artifact we refer to as spot swapping. Evidence for spot swapping is shown in Figure 1 in a tissue sample from postmortem human brain profiled as part of spatialLIBD, a project aimed at defining the spatial topography of gene expression in the six-layered human dorsolateral prefrontal cortex (DLPFC)^3^. Specifically, Figure 1a shows that UMI counts at background spots (which are zero in the absence of contamination) are high compared with counts in tissue spots; and the counts decrease with increasing distance from the tissue (Figure 1b). Figure 1c shows the distribution of UMI counts for 50 genes in a tissue region, a nearby background region, and a distant background region. As a result of expression similarity between the tissue and nearby background, tissue and background spots are not easily distinguished (Figure 1d). This is emphasized again in Figure 1f, where spots on the slide are colored by membership in the graph-based clusters shown in Figure 1e. Supplementary Figures 2-5 show similar results from 16 additional datasets; and Supplementary Table 1 shows that the proportion of UMI counts in background spots ranges between 5% and 20% in most datasets.

**Figure 2:**
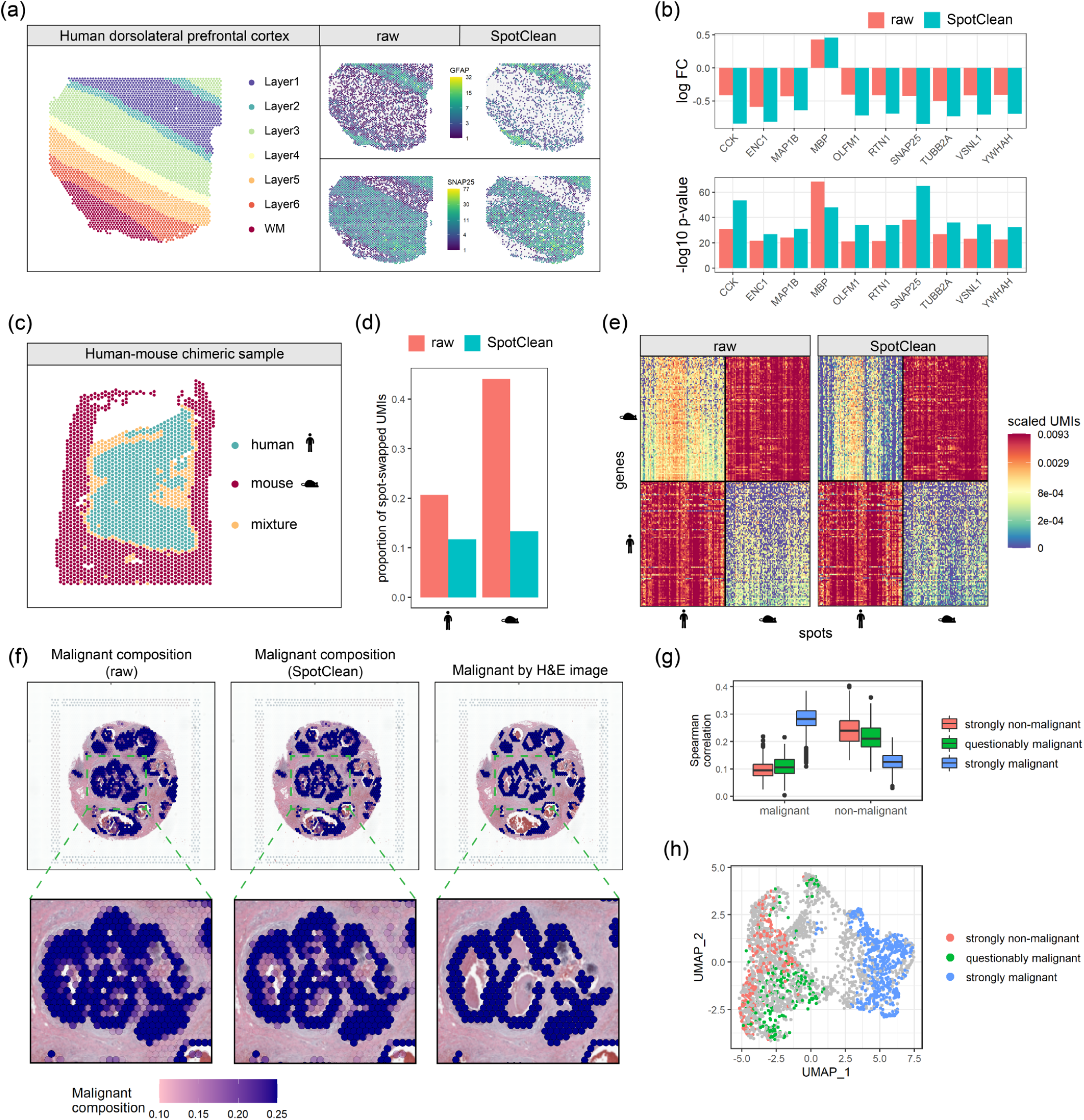
Data from the spatialLIBD study, sample LIBD_151507 (panels a and b); the chimeric experiment, sample HM-1 (panels c-e); and a human breast cancer study, sample human_breast_2 (panels f-h). (a) Known annotation of different layers of the human dorsolateral prefrontal cortex (left); layer-specific marker gene expression in the raw (middle) and SpotClean decontaminated (right) data show that SpotClean provides improved specificity of marker gene expression for GFAP, a marker for WM and Layer1, and for SNAP25, a neuronal marker up-regulated in Layer2-Layer6. (b) An analysis of genes known to be differentially expressed (DE) between WM and Layer6 in raw and SpotClean decontaminated data shows that SpotClean results in increased fold-changes and smaller p-values for the majority of known DE genes. (c) Species annotation of sample HM-1, a chimeric tissue of human skin and mouse duodenum. Spots annotated as mixtures were removed prior to calculating the summaries in panels (d) and (e) in an effort to ensure that the effects shown are not due to spots containing a mixture of the two species. (d) The proportion of spot-swapped UMI counts from all human genes (human-specific UMIs in background or mouse spots) are shown left for raw (salmon) and SpotClean decontaminated (turquoise) data; the proportion of spot-swapped UMI counts from all mouse genes (mouse-specific UMIs in background or human spots) are shown right. Note that there may be spot swapped UMIs within species (e.g. reads from human spot *t* bound by probes at human spot *t’*), but they cannot be identified in this experiment. (e) Scaled expression (UMIs are scaled so that each row sums to 1) for the top 100 human genes and top 100 mouse genes in the top 100 human spots and top 100 mouse spots. The top 100 human or mouse genes (spots) are those genes (spots) with highest total UMI counts. Data decontaminated via SpotClean shows reduced expression of human genes in mouse tissue, with no reduction in human tissue; and vice versa. (f) Malignant spot composition as estimated via SPOTlight is shown for the raw data (upper left) and SpotClean decontaminated data (upper middle). The raw data identifies many spots as malignant whereas the SpotClean decontaminated data more closely resembles the annotations derived from the H&E image (upper right). The inserts highlighted in the upper panel are shown in the lower panel. (g) Spearman correlations between average expression in the malignant scRNA-seq cells and spot-specific expression were calculated. Boxplots of correlations are shown for 265 strongly non-malignant spots, 216 questionably malignant spots (spots labelled malignant in the raw data, but not the SpotClean decontaminated data), and 546 strongly malignant spots. Correlations with non-malignant scRNA-seq cells are also shown. The correlations show that expression in the questionably malignant spots more closely resembles that in non-malignant cells suggesting that the malignant classification in the raw data at these spots is likely false due to spot swapping. (h) The UMAP plot further demonstrates that the questionably malignant spots are likely false positives as their expression more closely resembles that at non-malignant spots.

Figure 1, Supplementary Figures 2-5, and Supplementary Table 1 demonstrate that spot swapping occurs from tissue to background, but evaluating the extent of spot swapping from tissue spot to tissue spot is more challenging. While the SpotClean model provides an estimate (Supplementary Table 2), we also consider tissue-specific marker genes identified in the spatialLIBD project. In the absence of spot swapping, expression for a layer-specific marker should be high within that layer, and low (or off) in other layers. When spot swapping occurs, marker expression is relatively high in nearby layers. This is evident with GFAP, for example, a marker known to be up-regulated in white matter (WM) and in the first annotated layer of the DLPFC (Layer1). Supplementary Figure 6 shows high expression of GFAP in WM and Layer1 spots, as expected, but also relatively high expression in tissue spots adjacent to WM and Layer1, with GFAP expression decreasing as distance from WM (or Layer1) increases. While it is possible that some increase in marker expression in adjacent tissue spots may be due to the presence of WM (or Layer1) cells at those spots, we note that the rate of expression decay into the background spots (where no cells are present) is similar to the rate of decay into adjacent tissue regions. Consequently, the possible presence of WM (or Layer1) cells in adjacent tissue spots is not sufficient to fully explain the observed expression pattern. Similar results are shown for a WM marker, MOBP (Supplementary Figure 6), as well as 13 additional markers (Supplementary Figure 7).

To more directly quantify the extent of spot swapping, we conducted chimeric experiments where human and mouse tissues were placed contiguously during sample preparation. For each experiment, we annotated the H&E images to identify species-specific regions, and we calculated the proportion of spot-swapped reads (mouse-specific reads in human spots, human-specific reads in mouse spots, and reads in background spots). This is a lower bound on the <u>p</u>roportion of <u>s</u>pot-<u>s</u>wapped reads (LPSS) as it does not account for spot swapping within species (e.g. reads from human spot *t* bound by probes at human spot *t’*); LPSS ranges between 26-37% in these experiments (Supplementary Table 1). Taken together, results from a comparison of tissue and background expression (Figure 1 and Supplementary Figures 2-5), analysis of marker genes (Supplementary Figures 6-7), and the chimeric experiment (Supplementary Table 1 and Supplementary Figure 8) demonstrate that spot swapping affects UMI counts in ST experiments. This nuisance variability decreases the power and precision of downstream analyses (Figure 2b, Figure 2f-h, Supplementary Figure 9).

The statistical methods developed to adjust for known sources of contamination in RNA-seq experiments^4,5^ do not accommodate the spatial dependence inherent in spot swapping, and, consequently, are not sufficient in this setting (Supplementary Section S1). To adjust for the effects of spot swapping in ST experiments, we developed SpotClean. The approach is implemented in the R package *R/spotClean*. SpotClean was evaluated on simulated and case study data. In SimI, contaminated counts are generated assuming that local contamination follows a Gaussian kernel; SimII-IV relax the Gaussian assumption. In SimV, contaminated counts are simulated for genes having average expression that varies systematically across the slide. Supplementary Tables 3-6, which show the mean squared error (MSE) between true and decontaminated gene expression in simulated datasets, indicate that SpotClean provides better estimates of expression; and Supplementary Figure 10 demonstrates that SpotClean expression estimates lead to increased precision for identifying spatially varying genes.

The benefits of SpotClean on downstream analyses are also illustrated in case study data. Specifically, SpotClean increases the specificity of marker gene expression, increases the power for identifying DE genes, and improves the accuracy of spot annotations. Figure 2a shows that SpotClean improves the specificity of GFAP in the spatialLIBD data by maintaining expression levels in WM and Layer1 and reducing spurious expression in the other layers. Supplementary Figure 11 shows similar results for the 15 markers shown in Supplementary Figure 7. Figure 2b and Supplementary Figure 9 consider genes known to be differentially expressed (DE) between WM and Layer6 in raw and SpotClean decontaminated data; SpotClean results in increased fold-changes and smaller p-values for known DE genes. The chimeric datasets provide additional examples. In particular, Figure 2d shows that SpotClean reduces the proportion of spot-swapped UMI counts in the chimeric datasets. Similar results are shown in Figure 2e where we consider expression for human-specific and mouse-specific genes at human-specific and mouse-specific spots. Data decontaminated via SpotClean shows reduced expression of human genes in mouse tissue, with no reduction in human tissue, and vice versa.

There is considerable interest in applying spatial transcriptomics to personalized medicine, such as molecular profiling of patient tumor biopsies to guide diagnosis and precision therapy. SpotClean demonstrates substantial advantage in such applications where accurate spot annotation is crucial. Figure 2f shows a human breast cancer sample (ductal carcinoma), where the diagnosis and extent and invasiveness of tumor is typically estimated through evaluation of an H&E image by a pathologist. Spatial transcriptomics can provide additional information including identifying subtle collections of malignant cells, but accurate spot annotation is required for this information to be useful in clinical practice, and especially so as not to overcall tumor burden. Figure 2f shows spots annotated using SpotClean data versus spots annotated using data that has not been decontaminated via SpotClean. The non-decontaminated data misidentifies many spots as malignant including those containing benign inflammatory cells surrounding the tumor whereas the SpotClean decontaminated data more closely resembles identification of malignant cells on the H&E image. Figure 2g-h show that without SpotClean, over 13% of the spots labelled malignant in the raw data are likely false calls due to spot swapping.

Spatial transcriptomics provides unprecedented opportunity to address biological questions and enhance patient care, but artifacts induced by spot swapping must be adjusted for to ensure that maximal information is obtained from these powerful experiments. SpotClean provides for more accurate estimates of expression, thereby increasing the power and precision of downstream analyses.

## DATA AVAILABILITY

Raw sequence data for the 3 human-mouse chimeric experiments are available at GEO (accession number: GSE178221). Links to 14 public spatial transcriptomics datasets are available in Supplementary Table 7. The human breast cancer single-cell RNA-seq data from Chung *et al*.^6^ is available at GEO (accession number: GSE75688).

## CODE AVAILABILITY

The R package *SpotClean* is available at https://github.com/zijianni/SpotClean and will be submitted to Bioconductor. Codes for simulation and real data analyses as well as processed data can be found at https://github.com/zijianni/codes_for_SpotClean_paper.

## ACKNOWLEDGMENTS

This work was supported by NIH GM102756 and NIH UL1TR002373. The authors thank the University of Wisconsin Translational Research Initiatives in Pathology (TRIP) laboratory for assistance with sample preparation (P30 CA014520 and S10 OD023526) and the University of Wisconsin Biotechnology Center DNA Sequencing Facility for providing RNA sequencing facilities and services.

## AUTHOR CONTRIBUTIONS

Z.N. discovered the spot swapping artifact. Z.N. and C.K. designed the research and wrote the first version of the manuscript. Z.N., C.K., and M.N. developed the SpotClean method. A.P. and R.H. designed the chimeric samples and conducted the chimeric experiments. Z.N. and S.C. conducted simulations and quality control evaluations. Z.N., S.C. and C.K. built and tested the R package. All authors contributed to writing the manuscript.

## ONLINE METHODS

### Versions

The following software and packages were used in the analysis: R-4.0.2; R/SpotClean-0.99.0; R/SoupX-1.5.0; R/celda-1.5.11; R/Seurat-3.2.2; R/scran-1.17.20; R/SPOTlight-0.1.7; R/reticulate-1.16; Python-3.7.4; Python/spatialde-1.1.3; FastQC-0.11.7; MultiQC-1.9; Space Ranger-1.2.2; Loupe Browser-4.2.0.

### SpotClean

Let *K* be the total number of spots, *G* be the set of genes, *I*_*t*_ be the set of tissue spots with cardinality |*I*_*t*_| = *K*_*t*_, and *I*_*b*_ be the set of background spots with cardinality |*I*_*b*_| = *K*_*b*_ where *K*_*t*_ + *K*_*b*_ = *K*. The true (i.e., uncontaminated) UMI counts are given by 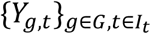 and observed counts by 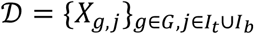. As our interest here is to characterize the extent of spot swapping, we introduce the missing variable *B*_*g,t,j*_ to be the UMI count for gene *g* leaving tissue spot *t* and binding to tissue (or background) spot *j*. Likewise we define *S*_*g,t*_ to be the UMI count arising from gene *g* in tissue spot *t* that remain at that spot and thus are not subject to bleeding. We decompose *Y*_*g,t*_ into a sum: *Y*_*g,t*_ = *S*_*g,t*_ + *B*_*g,t*_, where 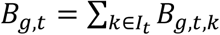 counts all bleed-outs from spot *t* to other spots *k* ≠ *t*. Extending notation, we set *Y*_*g,b*_ = *S*_*g,b*_ = *B*_*g,b*_ = 0 for background spots *b* ∈ *I*_*b*_ since background spots do not express mRNA. With these missing variables defined, we note that the measured count *X*_*g,j*_ = *S*_*g,j*_ + *R*_*g,j*_ where 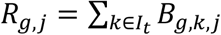 represents UMI counts received at spot *j* due to spot swapping. We leverage this missing-data formulation by flexibly modeling the component counts with independent Poisson distributions, which are known to be effective for UMI counts^7^.

For a collection of spot and gene-specific parameters, as well as global parameters controlling the swapping rates, we parameterize the distributions as: *S*_*g,t*_ ∼ Poisson(*μ*_*g,t*_ (1 − *r*_*β*_)) and 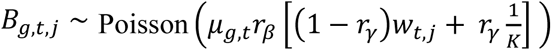 where *r*_*β*_ is the bleeding rate; *r*_*γ*_ is a distal and 1 − *r*_*γ*_ is a proximal contamination rate. By taking the global bleeding rate *r*_*β*_ ∈ [0,1], it follows that the uncontaminated counts follow: *Y*_*g,t*_ ∼ Poisson(*μ*_*g,t*_) for target parameters *μ*_*g,t*_ whose estimates constitute statistical estimates of the uncontaminated counts. Likewise for measured counts, *X*_*g,j*_ ∼ Poisson(*η*_*g,j*_), for induced gene and spot parameters. We define *w*_*t,j*_ by a weighted Gaussian kernel: *w*_*t,j*_ = *Kd*_*t,j*_, *σ*/ ∑_*j*′_ *Kd*_*t,j*′_, *σ* where *d*_*t,j*_ is the physical Euclidean distance between spots *t* and *j* measured in pixels in the slide image, σ is the kernel bandwidth, and 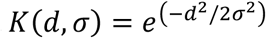 is a Gaussian kernel^8^.

### Parameter estimation

Plug-in estimates obtained by minimizing the residual sum of squares (RSS) between observed total counts and their expected values are used to estimate *r*_*β*_, *r*_*γ*_, and *σ* Specifically,

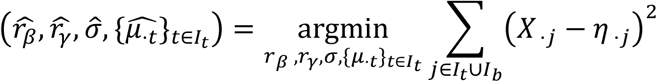

where *X*_*j*_, *η*_.*j*_, *μ*_*j*_ are the summations of *X*_*g,j*_, *η*_*g,j*_, *μ*_*g,j*_ among all genes, respectively. To reduce computational complexity, 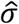 is taken as the minimum RSS calculated over a grid of candidate values. Explicit gradients are calculated for *r*_*β*_ and *r*_*γ*_ and estimates are obtained by L-BFGS-B gradient descent^9^. Details are provided in Supplementary Section S2. Since this optimization problem is not necessarily convex, it is important to choose appropriate initial values. For the initial values 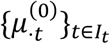 of 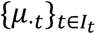, we use the observed total UMI counts 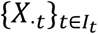 in tissue spots and scale them up so that they sum to the total UMIs in the data. The initial bleeding rate, 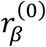, is the average expression in background spots divided by the average expression in all spots; and the initial distal contamination rate, 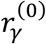, is defined by average expression in the 25^th^-50^th^ percentile of all background spots divided by average expression in all background spots.

With estimates 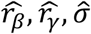 of the global parameters, true expression levels 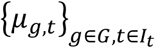 are readily estimated using an expectation-maximization (EM) algorithm^10^. Details are provided in Supplementary Section S3. For the initial values of true expressions 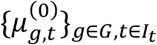, we use the observed UMI counts 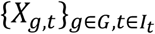 and scale up each gene so that their summations are equal to the gene summations in all spots.

### Estimation of spot-level contamination rate

For tissue spot *t*, let *c*_*t*_ be the proportion of contaminated UMIs from total observed UMIs. We estimate *c*_*t*_ using the estimated contamination received in *t* over its estimated contaminated total counts from model fitting: 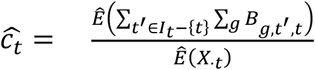. Validation of this estimate is provided in Supplementary Figure 12.

### Analysis of publicly available case study datasets

We downloaded UMI count matrices for 14 publicly available datasets, of which 12 came from 10x Visium and 2 came from Slide-seqV2^2^; links are provided in Supplementary Table 7. For each Visium dataset considered, the count matrix was normalized via scran^11^, following the Seurat^12^ pipeline for dimension reduction, clustering, and visualization. Seurat functions *FindVariableFeatures(nfeatures = 4000), ScaleData(), RunPCA(), RunUMAP(), FindNeighbors()*, and *FindClusters()* were applied under default settings. For each Slide-seqV2 dataset, we inspected total UMI counts of all spatial barcodes in the raw count matrix.

### Application of SoupX, DecontX, and SpotClean

Default parameters were used for SpotClean and DecontX. Since SoupX requires manual input of clusters, we first applied the Seurat pipeline on the raw tissue UMI count matrix to get cluster labels, with functions *NormalizeData(), FindVariableFeatures(), ScaleData(), RunPCA(), FindNeighbors(), FindClusters()* applied under default settings. Parameters for SoupX (*soupRange* in *estimateSoup(), tfidfMin* and *soupQuantile* in *autoEstCont()*) were manually tuned when the default settings failed. Some datasets did not run even after parameter tuning; results from these datasets are marked as NA. SpotClean decontaminates genes with average expression above 1, high variance as determined by Seurat’s *FindVariableFeatures()* function, or both. All methods were applied to these same set of genes. In the simulated data, we force all methods to decontaminate all genes since there are relatively few (1000 or 3000 genes depending on the simulation).

### Identification of marker genes and DE genes

The spatialLIBD project presented in Maynard *et al*.^3^ consists of spatial expression in the six-layered dorsolateral prefrontal cortex (DLPFC). The authors identified a number of marker genes for distinct layers of the DLPFC. In addition to these, we also considered marker genes from a single-cell RNA-seq study of Alzheimer’s disease^13^where markers differentiating between known cell types were identified. The markers shown here were selected from these papers if they were highly expressed (in the upper 25^th^ percentile) in the spatialLIBD datasets. We also evaluate the genes reported as DE between WM and Layer6 in Maynard *et al*.^3^. We filtered their list of DE genes and considered those genes having FDR<=10^−4^. From those, we chose the top 100 highest expressors in the raw data, sorted by fold change, and selected the top 10 for each dataset. For the DE analysis, raw and decontaminated tissue matrices were normalized using scran^11^; for each gene, p-values were obtained from a two-sample two-sided t-test between the 354 spots in WM and the 486 spots in Layer6. Summary statistics for the tests in Figure 2b are reported in Supplementary Tables 8-9.

### Human-mouse chimeric experiment

Fresh sections of normal human skin tissue were obtained with consent during routine dermatologic surgery under University of Wisconsin School of Medicine and Public Health Institutional Review Board (Approval #2010-0367). On the same day, fresh mouse tissue was harvested. All mouse husbandry and experimental procedures were performed in accordance and compliance with policies approved by the University of Wisconsin Research Animals Research and Compliance committee (Protocol #M5131). Three mixed species tissue blocks were then prepared under cold conditions as follows and frozen over a bed of dry ice and stored at -80°C in optimal tissue cutting (OCT) medium until they were ready to use:

HM-1: Duodenum from a 10-week-old C57BL/6J mouse as casing to a 4 mm punch section “cylinder” of human skin

HM-2: Colon from a 10-week-old C57BL/6J mouse as casing to a 4 mm punch section “cylinder” of human skin

HM-3: Heart from a 10-week-old C57BL/6J mouse encasing a 4 mm punch section “cylinder” of human skin

### Visium Spatial Transcriptomics

The Visium Spatial Tissue Optimization Slide & Reagent kit (10X Genomics) was used to optimize permeabilization conditions for the chimeric tissue according to manufacturer’s protocol and yielded an optimal tissue permeabilization time of 12 minutes. The Visium Spatial Gene Expression Slide & Reagent kit (10X Genomics) was used to generate sequencing libraries. Sections were cut at 10 μm thickness and mounted onto Visium slide capture areas, stained with H&E, digitally imaged, and then permeabilized for library preparation. Sequencing libraries were prepared following the manufacturer’s protocol. Initial quality control of the libraries was by analysis of 2×150 MiSeq data for each sample. The libraries were then sequenced on a NovaSeq 6000 (Illumina), with 29 bases from read 1 and 101 from read 2, at a depth of 500k-600k reads per spot. The actual depth was 455652, 440024, 538709 reads per spot for sample HM-1, HM-2, HM-3, respectively.

### Alignment and pre-processing in the chimeric experiment

The sequencing quality of each sample was evaluated using FastQC^14^ and MultiQC^15^. All FastQ files passed quality control. Tissues were manually aligned using the Loupe Browser. Reads were aligned to the GRCh38+mm10 reference genome (refdata-gex-GRCh38-and-mm10-2020-A from https://support.10xgenomics.com/single-cell-gene-expression/software/downloads/latest) and gene expression was quantified using Space Ranger under default parameters. Following alignment, we considered only those reads labeled confidently mapped by SpaceRanger; confidently mapped reads are reads that map uniquely to a gene. We refer to a gene as a human gene if it has prefix GRCh38; a mouse gene has prefix mm10. UMI counts were normalized for differences in total counts across species by scaling total UMI counts in mouse to match total UMI counts in human. Genes having average expression <0.01 were removed.

### Human and mouse tissue spot annotation in the chimeric experiment

Tissue spots were labelled as human, mouse, or histopathological mixture based on visual inspection of the H&E images. A histopathological mixture spot is one with tissue contributions from both species that can be visually verified in the H&E stained image. A pure human or pure mouse spot was relabeled as a computational mixture spot if the spot label differed from the majority of UMIs. Specifically, a human (or mouse) spot was labelled as a computational mixture if the total UMI counts from mouse (human) exceeded the median of total UMI counts across all mouse spots (human spots). Both histopathological or computational mixture spots were removed prior to analyses in an effort to ensure that the effects shown are not due to spots containing a mixture of the two species.

### Lower bound on the proportion of spot swapped reads (LPSS)

Spot swapped reads include reads from one tissue spot binding background probes (tissue-to-background) as well as reads at one tissue spot binding probes at another tissue spot (tissue-to-tissue). It is not possible to directly measure tissue-to-tissue swapping in most cases. However, the chimeric experiment provides some insight into the extent of spot swapping tissue-to-tissue. We define LPSS in the chimeric experiment as the proportion of misclassified reads (mouse reads in human spots, human reads in mouse spots, and reads in background spots). This is a lower bound as it does not account for spot swapping within species (e.g. reads from human spot *t* bound by probes at human spot *t’*).

### Cell type decomposition of the human breast cancer data

For cell type decomposition, we applied SPOTlight^16^ to the Visium human breast cancer data (referred to here as human_breast_2; details on this data are provided in Supplementary Table 7). SPOTlight^16^ requires single-cell RNA-seq data to use as a reference; for this, we used the human breast cancer single-cell RNA-seq data from Chung *et al*.^6^ SPOTlight^16^ was applied to the raw data under default settings to estimate the cell type composition of every spot; SPOTlight^16^ was also applied to the SpotClean decontaminated data under default settings. Note that since tumor cell populations are heterogeneous, and spots contain multiple cells, most spots containing malignant cells will also contain non-malignant cells. Following clinical practice, we label a spot as malignant if there is any evidence of malignancy. Specifically, we annotate spots as malignant if the estimated malignant cell composition exceeds 10%, which corresponds to approximately 1 malignant cell in the spot since the estimated number of cells in a spot is approximately 10 in Visium data^16^. We further define non-malignant spots as “strongly non-malignant” if the non-malignant cell composition exceeds 95%, and “strongly malignant” if the malignant cell composition exceeds 30% in both raw and decontaminated data. “Questionably malignant” is used to refer to spots called malignant in the raw data, but not the SpotClean decontaminated data. Spearman correlations between the expression of each spot and the average expression of malignant cells in the reference single-cell data were calculated to measure the similarity of each spot group (strongly non-malignant, strongly malignant, or questionably malignant) to malignant cells; the same was done to measure similarity of each spot group to non-malignant cells. Boxplots in Figure 2g demonstrate the median, upper and lower quartile, range without outliers, and outlier values of the Spearman correlations for each group of spots using default plotting functions. The Seurat pipeline, as described previously, was applied under default settings to the decontaminated data to produce the UMAP plot. In the H&E image, tissue spots were labelled as malignant and non-malignant based on visual inspection.

### Simulations

SimI simulates the spot swapping effect to get contaminated UMI counts given an input dataset. Specifically, starting from an input UMI count matrix of real data, 3000 genes with highest total UMI counts were selected. Expression for these genes was scaled to target the same average UMI total counts (average taken over spots) across input datasets. Denote the resulting matrix by 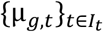. The bleeding rate *r*_*β*_ and distal contamination rate *r*_*γ*_ were estimated from the input data, using the same approach as described for obtaining initial values in SpotClean. The spot distances 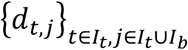 were calculated based on the spot coordinates in the H&E image of the input dataset; the contamination radius, *σ*, was set to 10; and the weights which describe the proportion of UMIs swapping locally from tissue spot *t* to any spot *j, w*_*t,j*_, is given by a Gaussian kernel. The expected contamination of gene *g* from tissue spot *tt* to spot *j* is then given by 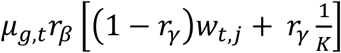. Summing contamination from all tissue spots to spot *j* and adding the UMIs that stay at *j, μ*_*g,j*_(1 − *r*_*β*_), gives the expected observed expression *η*_*g,j*_. Simulated counts for gene *g* in spot *j* are sampled from Poisson(*η*_*g,j*_).

Additional simulations are similar, but proximal contamination weights are not given by a Gaussian kernel. Rather, SimII, SimIII, and SimIV assume proximal contamination weights are given by a Linear, Laplace, and Cauchy kernel, respectively.

For SimV, starting from a UMI count matrix of real data, we select the top 5000 most highly expressed genes; any gene having average expression less than 0.1 is removed. SpatialDE^17^ is then applied using default settings; the top 500 highest expressed genes with q-value <=0.01 are identified as true spatially variable (SV) genes. For each SV gene, we simulate a matched non-SV gene by sampling independent Poisson counts parameterized by the average expression of the SV gene.

