## Supplementary figures, tables, and notes for "SpotClean adjusts for spot swapping in spatial transcriptomics data"

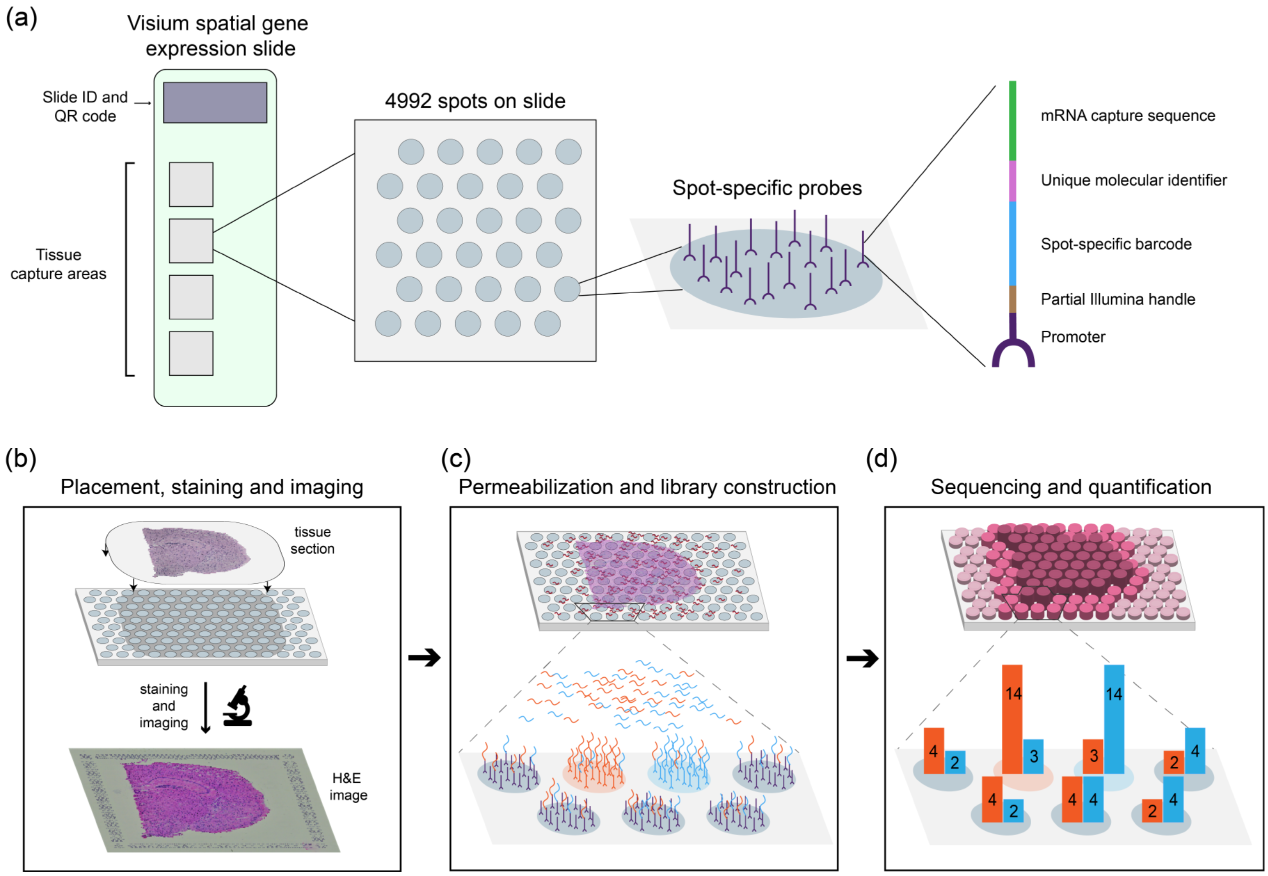


**Supplementary Figure 1:** Overview of the 10x Genomics Visium (10x)[1] spatial transcriptomics experiment. (a) The 10x Visium spatial gene expression slide contains four tissue capture sites for processing multiple tissue samples simultaneously. Each capture site contains 4992 spots, with each spot containing millions of spot-specific probes that bind mRNA. (b) Fresh frozen or FFPE tissue is sectioned, placed on a capture area, and imaged, typically via Hematoxylin and Eosin (H&E) staining. (c) Following imaging, the tissue is permeabilized to release mRNA. The lower panel shows five background spots (gray) and two tissue spots (orange and blue). Due to spot swapping, mRNAs from one tissue spot bind probes at other spots. (d) The bound mRNAs are released, processed, sequenced and quantified to give a gene-by-spot matrix of UMI counts. In this hypothetical example, due to spot swapping, UMI counts at each of the seven spots are a mixture of mRNAs from the two distinct tissue spots.

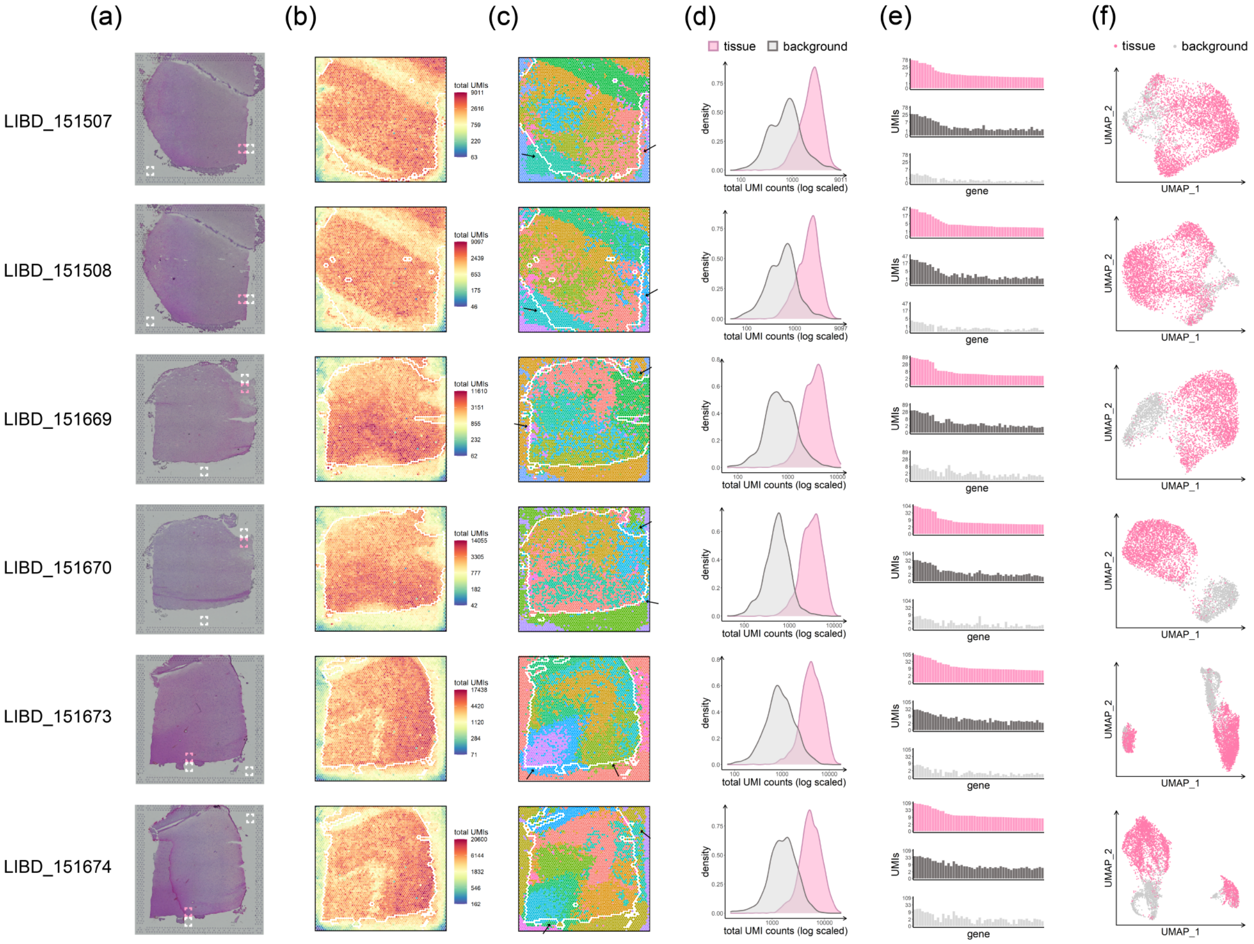


**Supplementary Figure 2:** Data from six human dorsolateral prefrontal cortex samples in the spatialLIBD project[2]. (a) H&E images for the six different samples. (b) UMI total counts in the background decrease with increasing distance from the tissue. (c) Spots on the slide are colored by their cluster membership via graph-based clustering (clusters not shown). Black arrows highlight areas of spot swapping. (d) UMI count densities for tissue and background spots show relatively high counts in the background. (e) Counts of the top 50 genes (genes with highest total UMI expression) from a select tissue region (upper), from a nearby background region (middle), and from a distant background region (bottom) show the similarity between expression in tissue spots and nearby background spots due to spot swapping from tissue to background, an effect that decreases as distance from the tissue increases. The tissue region and background regions used for each sample are highlighted in panel (a) in pink and white, respectively. (f) Spot similarities are visualized via UMAP plots; tissue and background spots are shown in pink and grey, respectively. There is considerable overlap of tissue and background spots in the UMAP plots. Tissue spots on the perimeter (shown in white in panels (b) and (c)) were removed prior to calculating the summaries in panels (d)-(f) in an effort to ensure that the effects shown are not due to spots on the tissue-background boundary.

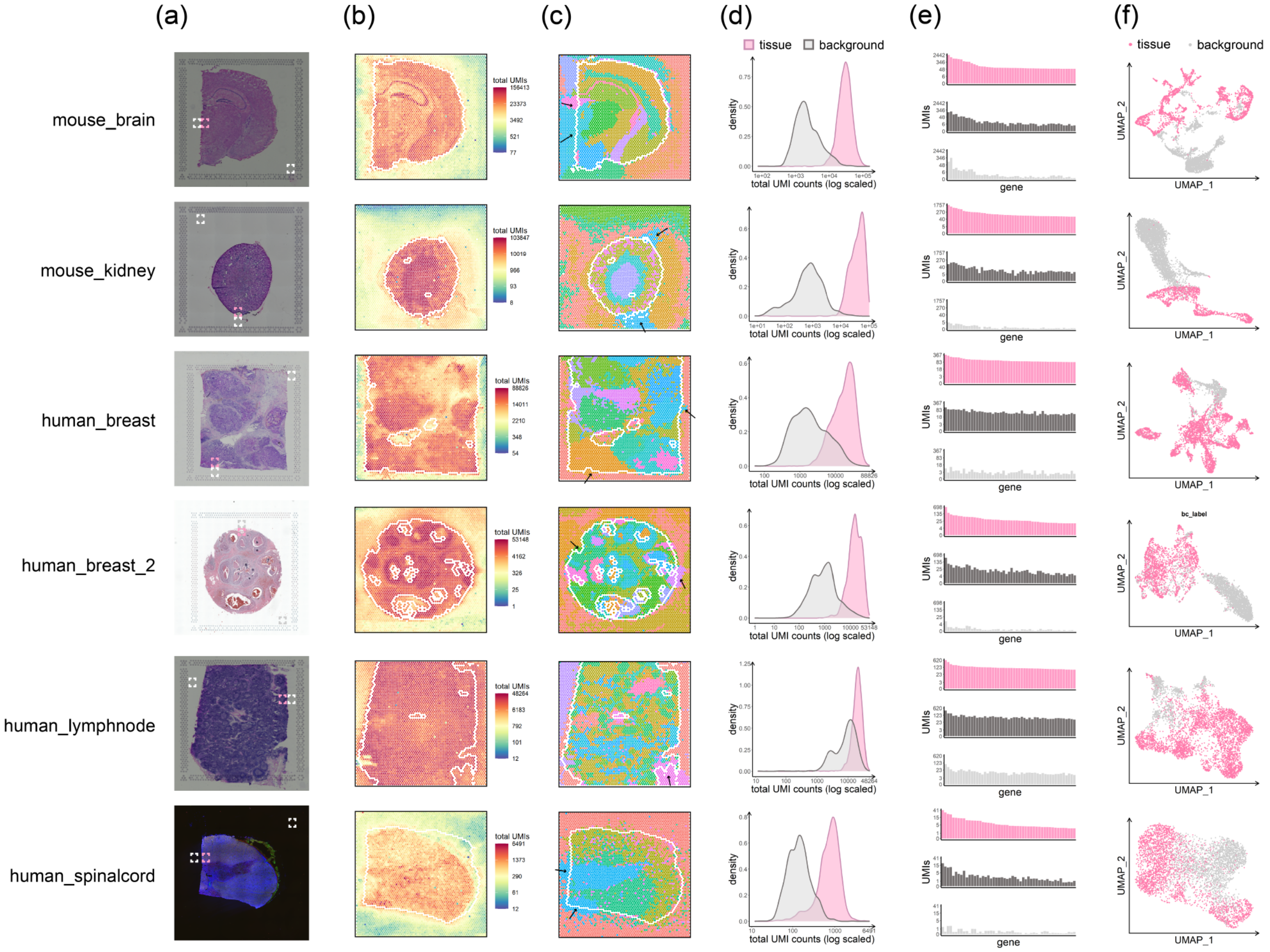


**Supplementary Figure 3:** Data from six publicly available 10x Genomics datasets. (a) H&E images for the six different samples. (b) UMI total counts in the background decrease with increasing distance from the tissue. (c) Spots on the slide are colored by their cluster membership via graph-based clustering (clusters not shown). Black arrows highlight areas of spot swapping. (d) UMI count densities for tissue and background spots show relatively high counts in the background. (e) Counts of the top 50 genes (genes with highest total UMI expression) from a select tissue region (upper), from a nearby background region (middle), and from a distant background region (bottom) show the similarity between expression in tissue spots and nearby background spots due to spot swapping from tissue to background, an effect that decreases as distance from the tissue increases. The tissue region and background regions used for each sample are highlighted in panel (a) in pink and white, respectively. (f) Spot similarities are visualized via UMAP plots; tissue and background spots are shown in pink and grey, respectively. There is considerable overlap of tissue and background spots in the UMAP plots. Tissue spots on the perimeter (shown in white in panels (b) and (c)) were removed prior to calculating the summaries in panels (d)-(f) in an effort to ensure that the effects shown are not due to spots on the tissue-background boundary.

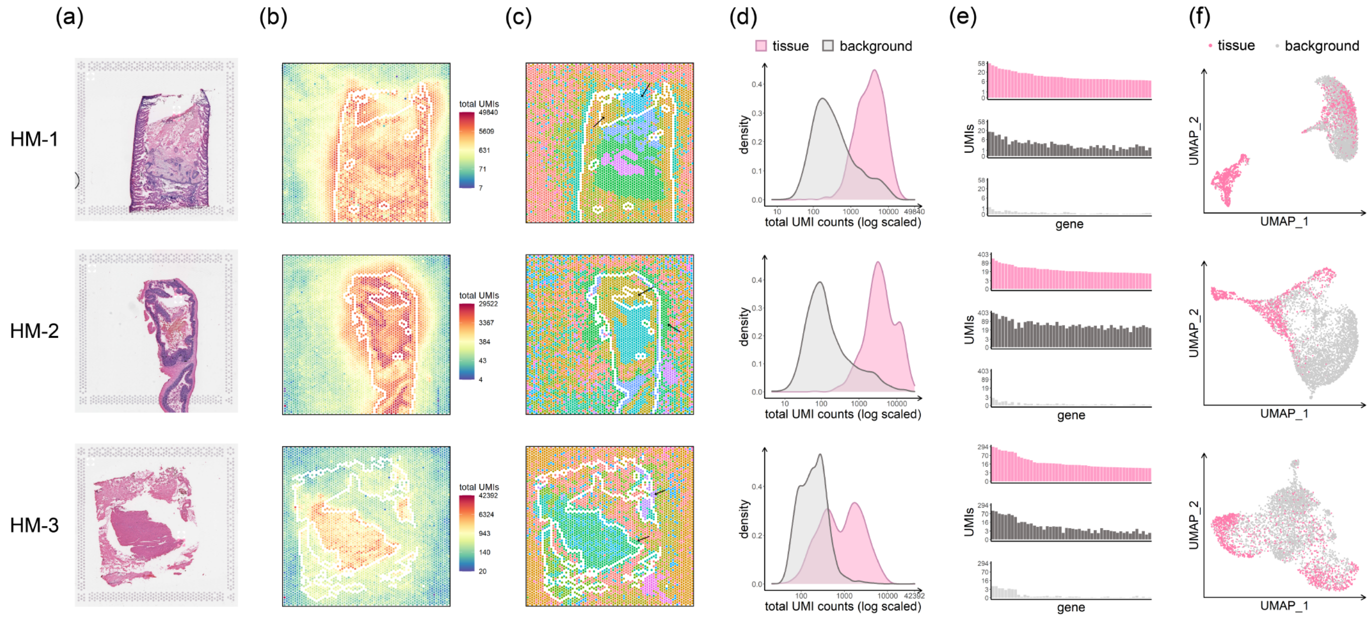


**Supplementary Figure 4:** Data from the three chimeric samples composed of human and mouse tissues. (a) H&E images for the three different samples. (b) UMI total counts in the background decrease with increasing distance from the tissue. (c) Spots on the slide are colored by their cluster membership via graph-based clustering (clusters not shown). Black arrows highlight areas of spot swapping. (d) UMI count densities for tissue and background spots show relatively high counts in the background. (e) Counts of the top 50 genes (genes with highest total UMI expression) from a select tissue region (upper), from a nearby background region (middle), and from a distant background region (bottom) show the similarity between expression in tissue spots and nearby background spots due to spot swapping from tissue to background, an effect that decreases as distance from the tissue increases. The tissue region and background regions used for each sample are highlighted in panel (a) in pink and white, respectively. (f) Spot similarities are visualized via UMAP plots; tissue and background spots are shown in pink and grey, respectively. There is considerable overlap of tissue and background spots in the UMAP plots. Tissue spots on the perimeter (shown in white in panels (b) and (c)) were removed prior to calculating the summaries in panels (d)-(f) in an effort to ensure that the effects shown are not due to spots on the tissue-background boundary.

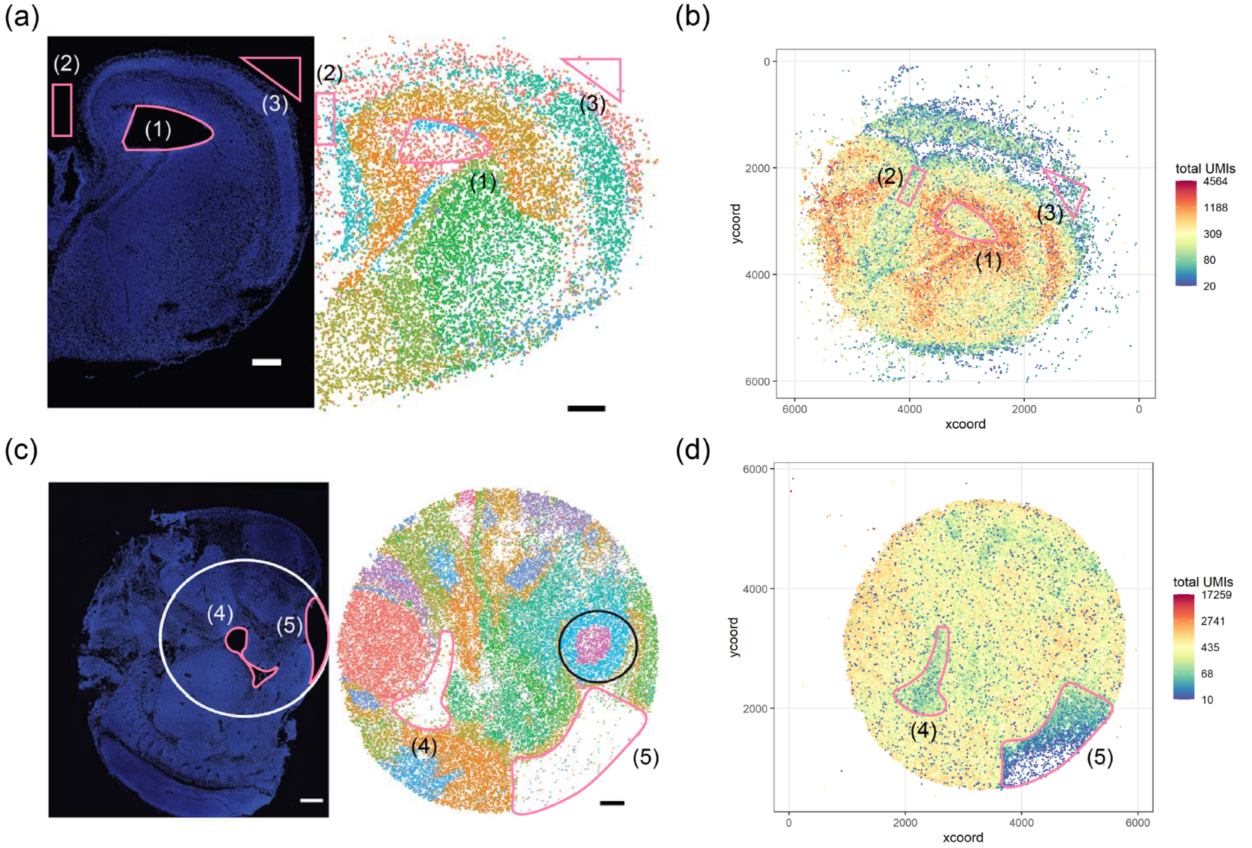


**Supplementary Figure 5:** Evidence of spot swapping in Slide-seqV2 data. (a) DAPI stain of E15 mouse brain (left) and Slide-seqV2 data of E15 brain with cluster labels (right) reported in Stickels *et al.*[3]. Three background regions identified from the DAPI image are shown in pink and labeled as (1), (2) and (3). The same regions are also identified in the graph-based clustering of beads. (b) Raw UMI counts data colored by total UMI counts for all spatial barcodes shows positive UMI counts detected in the three background regions. (c)-(d) are identical to (a)-(b), but for the E12.5 mouse embryo data. The black circle shown in (c) is part of the original image and not relevant here.

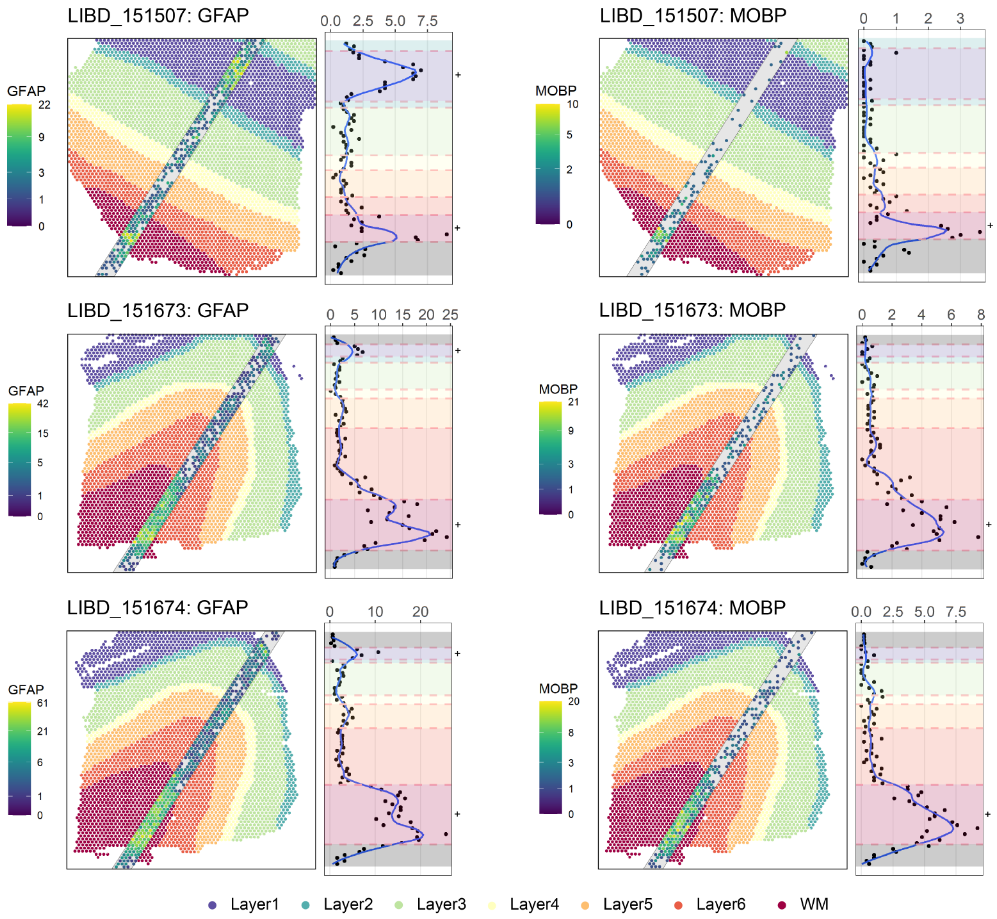


**Supplementary Figure 6:** The effect of spot swapping on layer-specific marker genes in three spatialLIBD datasets. The upper left panel shows the annotated LIBD_151507 sample and a stripe with width equal to five spots. Each spot in the stripe is colored by expression of GFAP, a marker for white matter (WM) and Layer1; the "+" denotes the regions where marker expression is expected to be high (here, WM and Layer 1). Average expression of each row in the stripe is shown in the right subpanel. The average is taken across the five spots for every row contained completely within a layer; for rows containing two layers, the average is taken across the three or four spots making up the major layer. When spot swapping occurs, marker expression is relatively high in nearby layers, as observed here. While it is possible that some increase in marker expression in adjacent tissue spots may be due to the presence of WM (or Layer1) cells at those spots, we note that the rate of expression decay into the background spots (where no cells are present) is similar to the rate of decay into adjacent tissue regions. Consequently, the possible presence of cells from a given layer in adjacent tissue spots outside that layer is not sufficient to fully explain the observed expression patterns shown here. The lower middle and lower left panels show the same plot for different tissue samples; the right panels show MOBP, another WM marker.

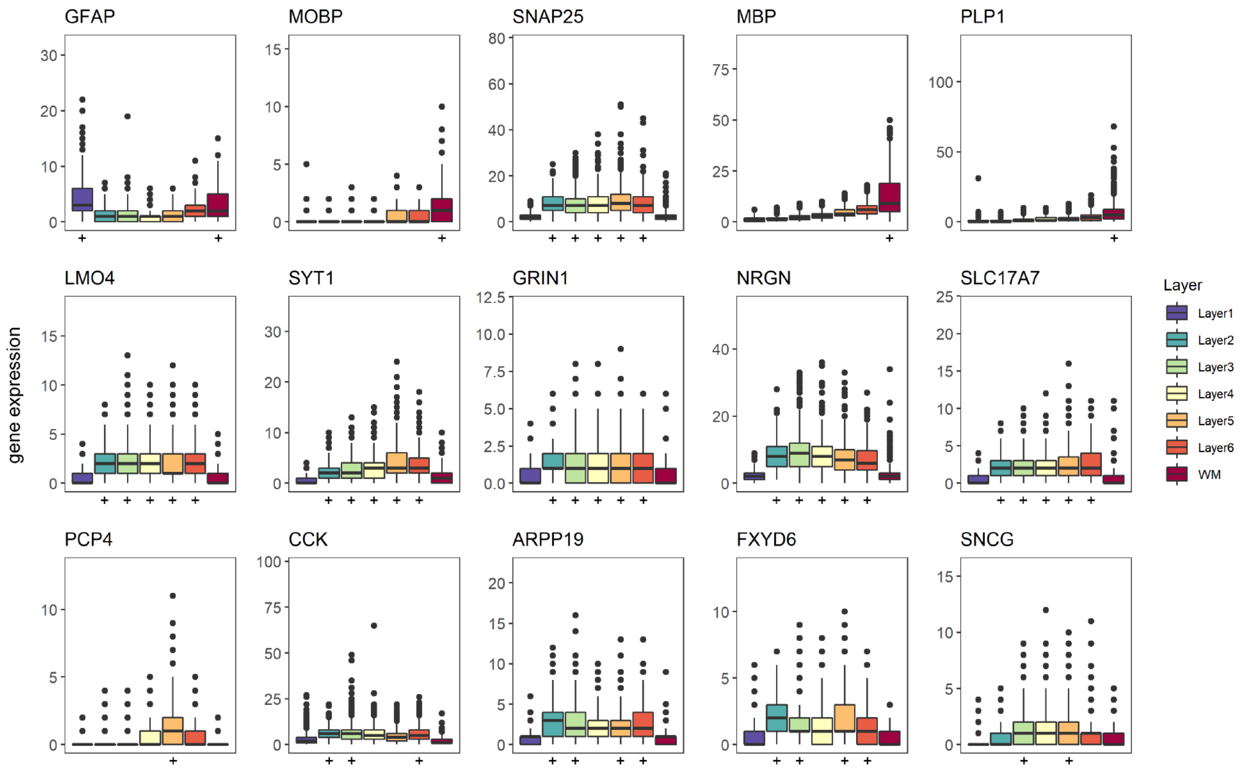


**Supplementary Figure 7:** Boxplots of expression within each layer in LIBD_151507 sample are shown for 15 layer-specific markers identified in the spatialLIBD project. The "+" denotes the layer(s) for which expression is expected to be high. In the absence of spot swapping, expression for a layer-specific marker should be high within that layer, and low (or off) in other layers. Observed expression is relatively high in adjacent tissue regions for most of the markers considered.

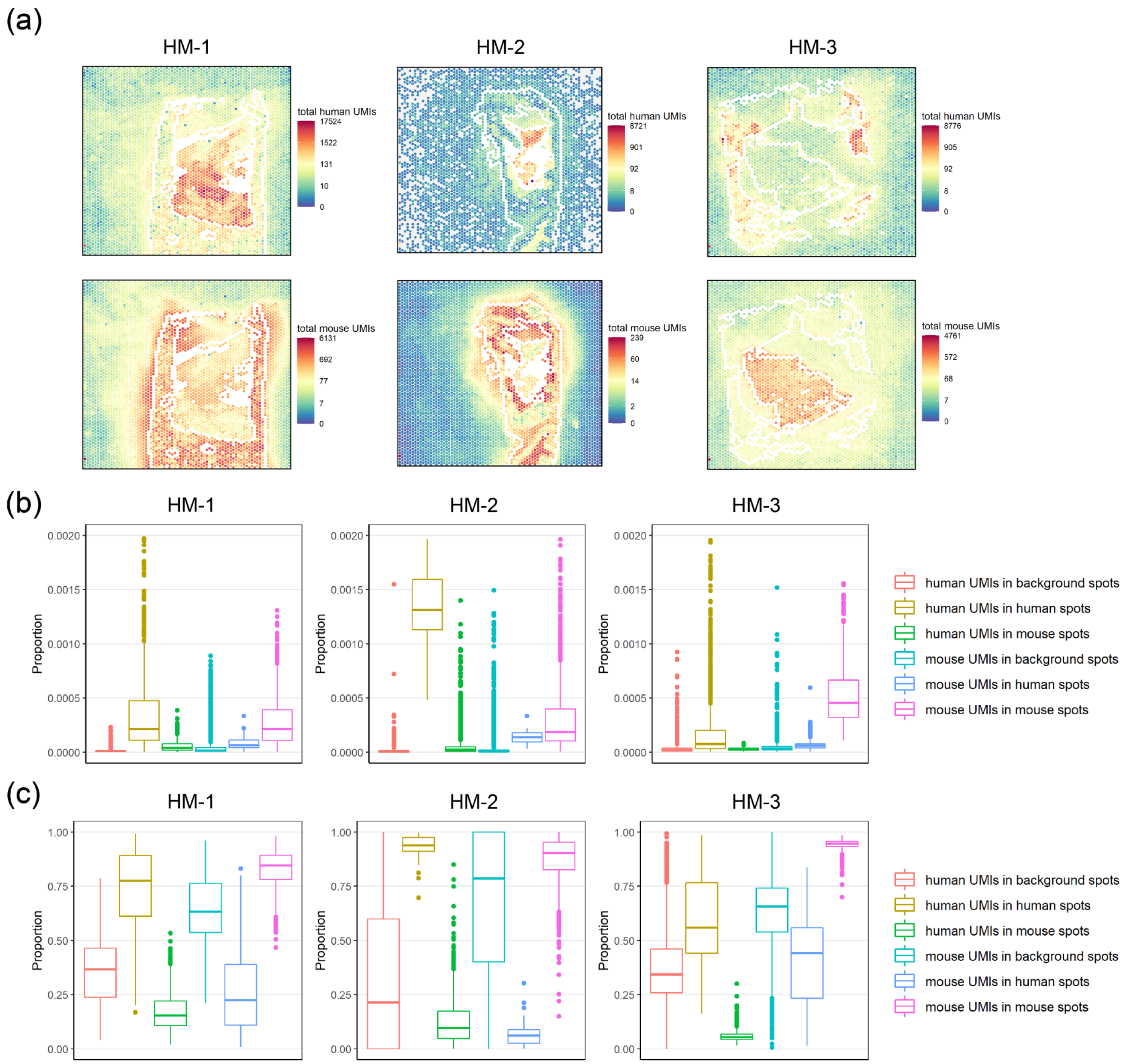


**Supplementary Figure 8**: Panel (a) shows total UMI counts in human-specific genes (upper) and mouse-specific genes (lower) for the three chimeric experiments. Panel (b) shows the proportion (out of total UMIs) of spot-swapped UMI counts (human-specific UMIs in background or mouse spots; mouse-specific UMIs in background or human spots). Also shown are the proportion of human-specific UMIs in human spots and mouse-specific UMIs in mouse spots. Note that there may be spot swapped reads in these latter proportions (e.g. reads from human spot $t$ bound by probes at human spot $t'$), but they cannot be identified in this experiment. Panel (c) shows spot-specific proportions. Tissue spots on the perimeter as well as spots annotated as mixtures (shown in white in panel (a)) were removed prior to calculating the summaries in panels (b) and (c) in an effort to ensure that the effects shown are not due to spots on the tissue-background boundary.

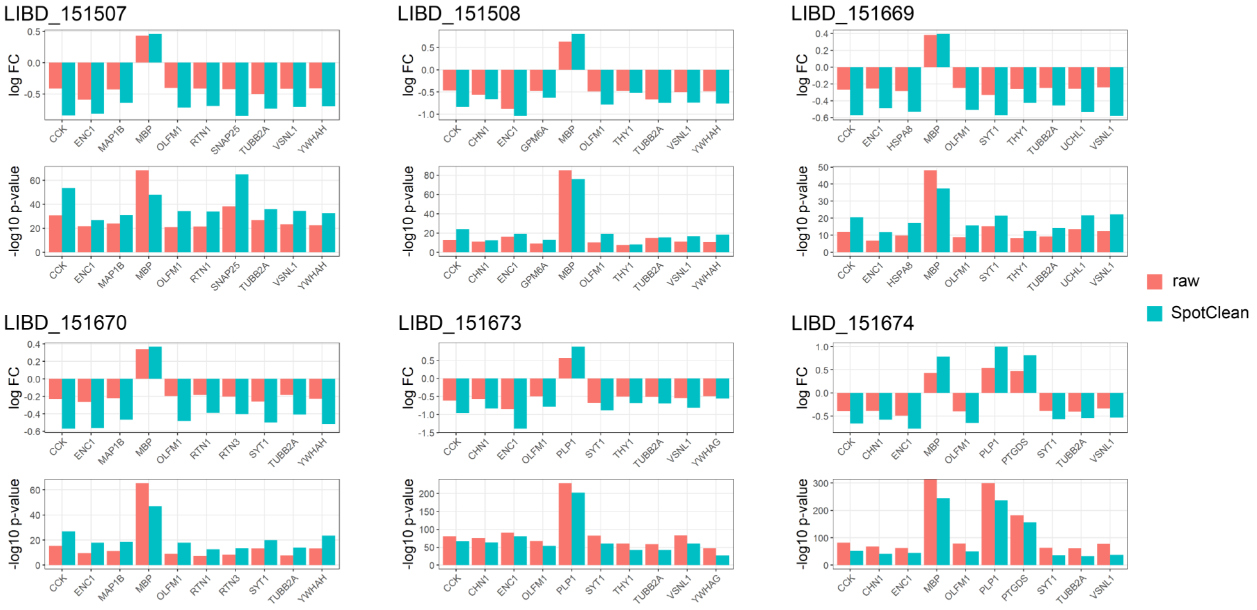


**Supplementary Figure 9**: Shown for the 6 spatialLIBD datasets are the fold changes and p-values for 10 genes known to be differentially expressed (DE) between WM and Layer6 for the raw data (salmon) and SpotClean processed data (turquoise). By reducing noise due to spot-swapped UMIs, SpotClean improves fold changes and p-values for the majority of known DE genes.

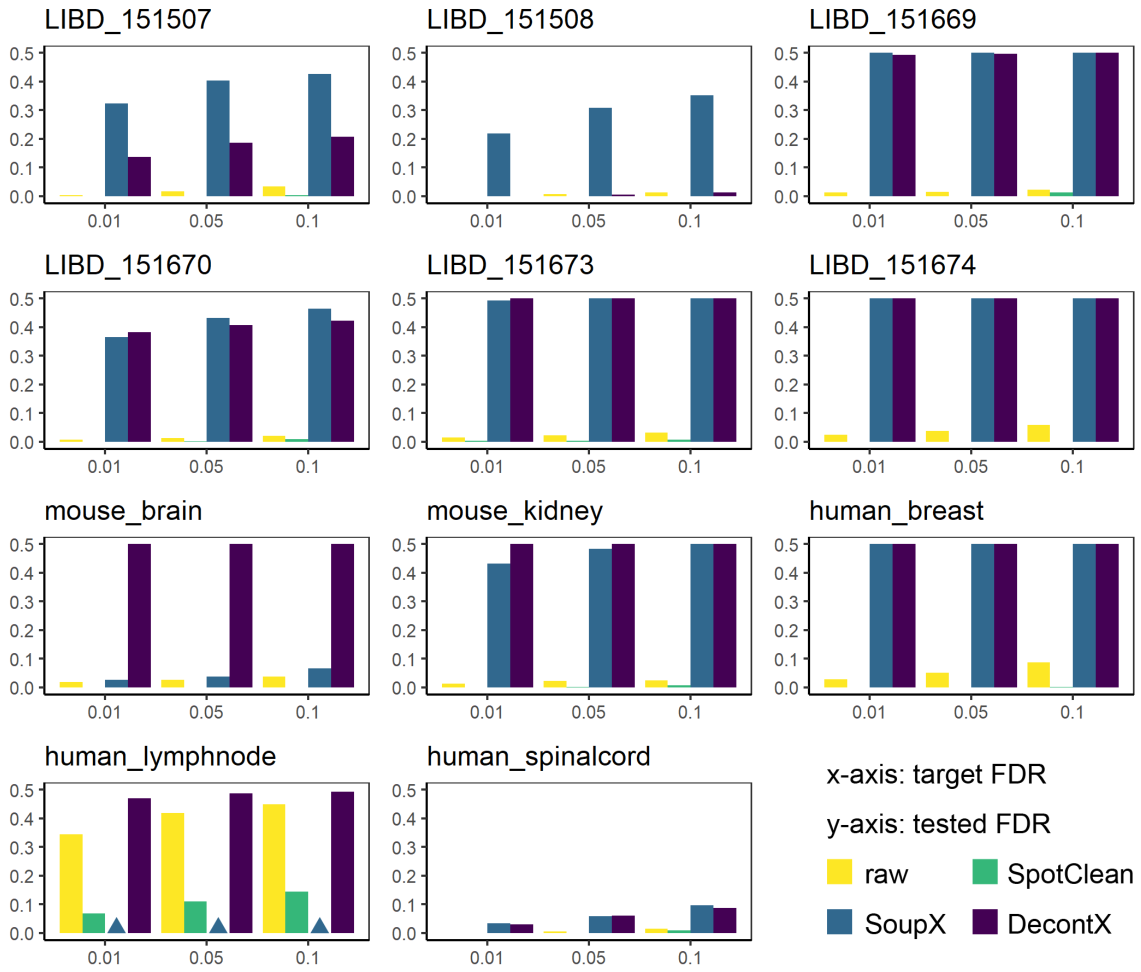


**Supplementary Figure 10**: SimV simulated data containing spatially varying (SV) genes was simulated for 11 datasets using input from the publicly available dataset indicated. SpatialDE[4] was applied to the simulated data following decontamination by SpotClean, SoupX[5], and DecontX[6]. Shown is the observed false discovery rate (FDR) for the raw data and each dataset following decontamination for three target FDR cut-offs. The upper bound on observed FDR is 0.5 since there are 50% simulated SV genes and 50% simulated non-SV genes. SoupX failed to run on the human_lymphnode data (NA shown as triangles). SoupX and DecontX have increased FDR for many datasets as they impose variability on EE genes during decontamination (Supplementary Section S1 and Supplementary Figure 13).

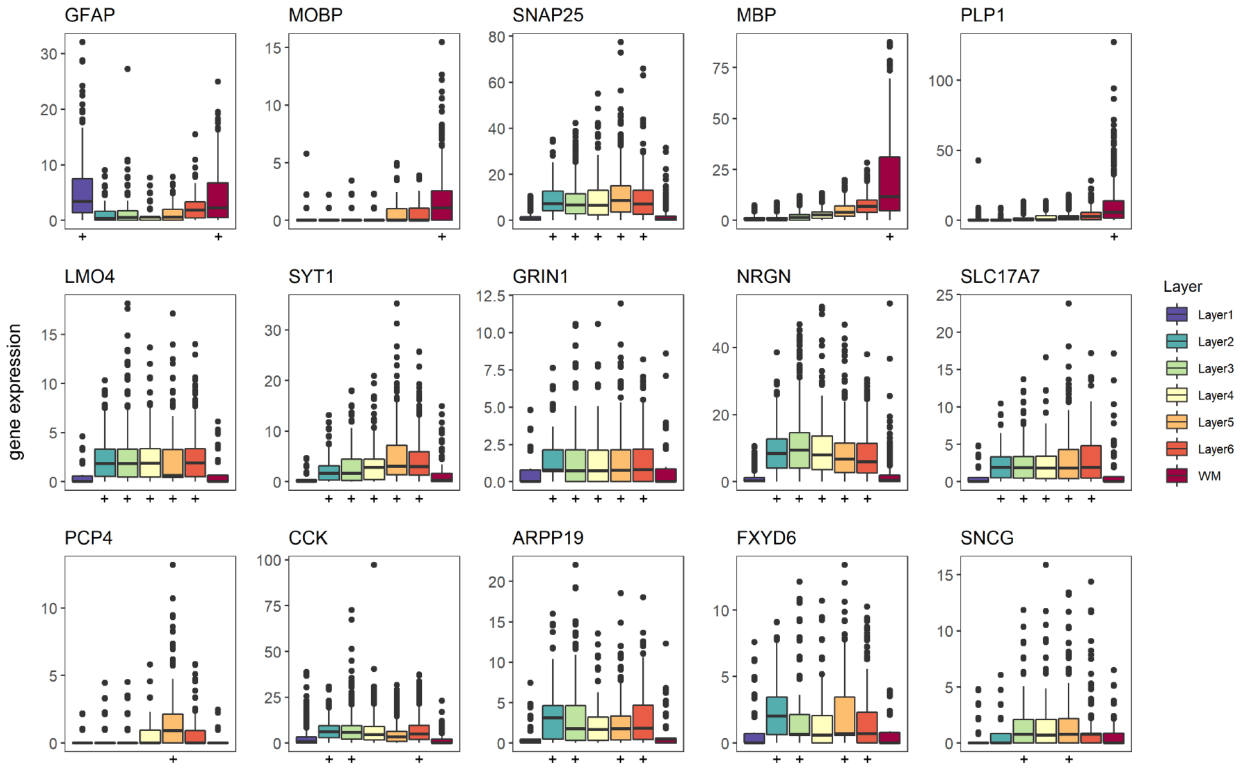


**Supplementary Figure 11:** Boxplots of expression estimates from SpotClean are shown within each layer for 15 layer-specific markers identified in the spatialLIBD project. The "+" denotes the layer(s) for which expression is expected to be high. Compared with Supplementary Figure 7, for most markers we observe increased expression in layers for which the marker is expected to be high, and reduced expression in adjacent tissue regions since SpotClean reduces the effect of spot-swapping.

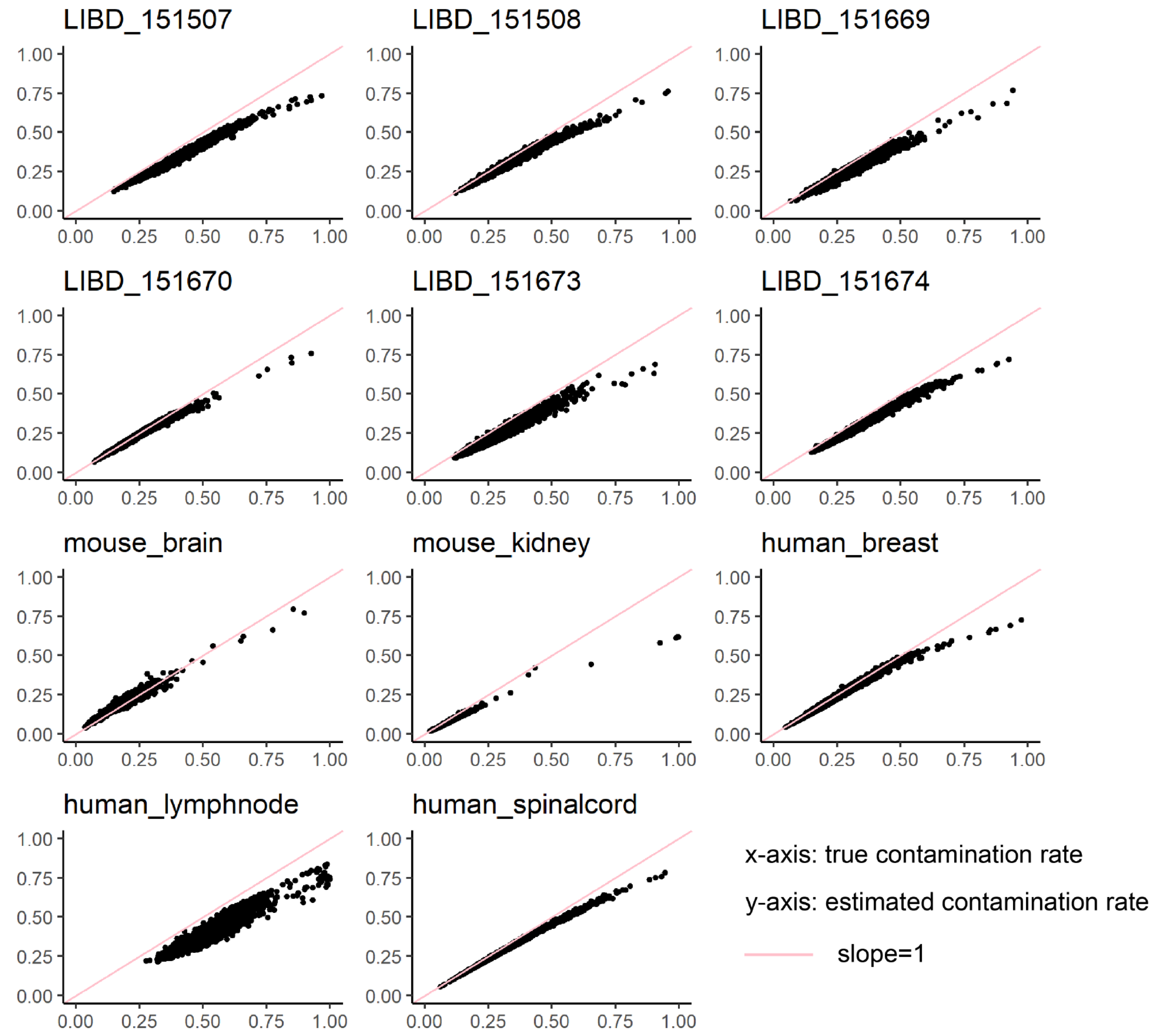


**Supplementary Figure 12**: Contamination rates estimated via SpotClean (y-axis) are plotted against true contamination rates (x-axis) for 11 datasets simulated using input from the dataset indicated. SpotClean provides reasonable estimates of the contamination rate for most simulated datasets and tends to be conservative when the true contamination rate is large.

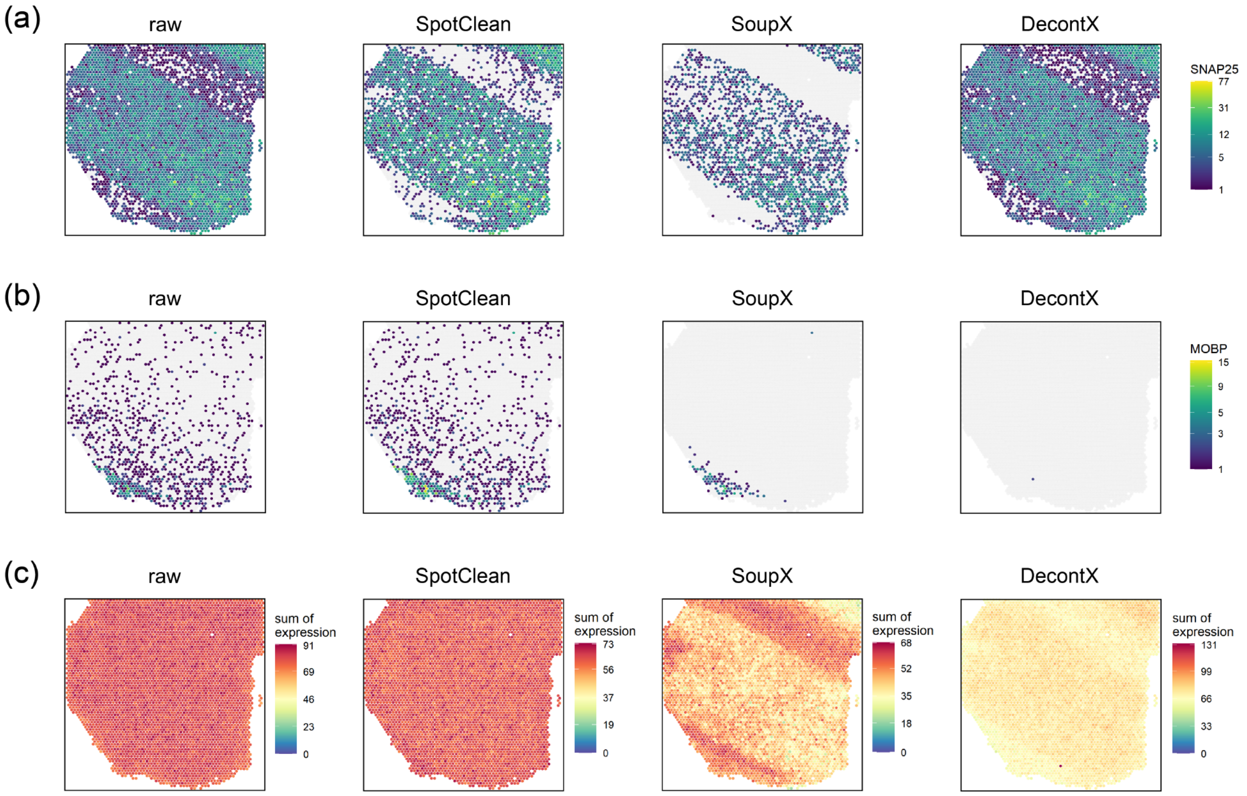


**Supplementary Figure 13:** Data from the spatialLIBD experiment, sample LIBD_151507. Data decontaminated by SpotClean (middle left), SoupX (middle right), and DecontX (right) for SNAP25 (a) and MOBP (b). The raw data is shown left. SoupX decontaminates SNAP25, but reduces expression even in Layer2-Layer6, where SNAP25 expression is expected to be high; DecontX imposes little change on this marker's expression. SoupX works well for MOBP, but DecontX removes almost all of the signal for this marker. Panel (c) shows results from SimV data generated using sample LIBD_151507; 500 spatially varying (SV) genes and 500 genes showing no change in expression across the slide were simulated (non-SV). Shown far left in panel (c) is summed expression for the 500 non-SV genes. To ensure that the summation is not dominated by a few highly expressed genes, gene-specific expression was scaled so that the maximum value of each gene equals 1. The same sum is shown for data decontaminated by SpotClean (middle left), SoupX (middle right), and DecontX (right). SoupX and DecontX impose artificial patterns upon non-SV genes.

| Dataset | Prop. UMIs in Background | Number of background spots | Normalized prop. UMIs in Background | LPSS |
|---|---|---|---|---|
| LIBD_151507 | 6.4% | 766 | 0.0084% | NA |
| LIBD_151508 | 4.2% | 606 | 0.0069% | NA |
| LIBD_151669 | 8.3% | 1329 | 0.0062% | NA |
| LIBD_151670 | 7.6% | 1494 | 0.0051% | NA |
| LIBD_151673 | 8.6% | 1353 | 0.0064% | NA |
| LIBD_151674 | 10.4% | 1319 | 0.0079% | NA |
| mouse_brain | 8.3% | 2290 | 0.0036% | NA |
| mouse_kidney | 7.7% | 3554 | 0.0022% | NA |
| human_breast | 4.3% | 1005 | 0.0043% | NA |
| human_lymphnode | 10.7% | 957 | 0.0112% | NA |
| human_spinalcord | 14.2% | 2180 | 0.0065% | NA |
| HM-1 | 20.8% | 2962 | 0.0070% | 32.6% |
| HM-2 | 17.6% | 3666 | 0.0048% | 26.0% |
| HM-3 | 26.9% | 3092 | 0.0087% | 36.6% |

**Supplementary Table 1**: The proportion of UMI counts in background spots, the number of background spots, the per-spot proportion of UMI counts in background spots, and the LPSS (which is only defined for the three chimeric datasets). The proportion of UMI counts in background spots serves as an underestimate of the proportion of spot-swapped UMI counts (since the proportion quantifies tissue-to-background swapping but does not account for tissue-to-tissue swapping). The proportion depends on tissue coverage since experiments with relatively few background spots will necessarily have a lower proportion of UMI counts in the background. Consequently, we also report the normalized proportion, which is the proportion of UMI counts in background spots divided by the number of background spots in each dataset. LPSS is defined in the chimeric experiment as the proportion of misclassified reads (mouse reads in human spots, human reads in mouse spots, and reads in background spots). This is a lower bound as it does not account for spot swapping within species (e.g. reads from mouse spot $t$ bound by probes at mouse spot $t'$).

| Dataset | Mean est. contamination rates | Median est. contamination rates |
|---|---|---|
| LIBD_151507 | 38.5% | 36.4% |
| LIBD_151508 | 39.3% | 37.2% |
| LIBD_151669 | 39.9% | 39.1% |
| LIBD_151670 | 36.8% | 35.5% |
| LIBD_151673 | 39.6% | 38.4% |
| LIBD_151674 | 41.4% | 39.9% |
| mouse_brain | 31.4% | 29.2% |
| mouse_kidney | 30.0% | 26.6% |
| human_breast | 40.7% | 37.6% |
| human_lymphnode | 44.4% | 42.3% |
| human_spinalcord | 38.8% | 35.7% |
| HM-1 | 40.1% | 37.6% |
| HM-2 | 41.5% | 36.8% |
| HM-3 | 48.2% | 48.7% |

**Supplementary Table 2**: Spot-swapped reads include reads at one tissue spot binding probes at another (tissue-to-tissue) as well as reads from one tissue spot binding background probes (tissue-to-background). While it is not possible to directly measure tissue-to-tissue swapping in most cases, the SpotClean model provides an estimate. The mean and median of spot-wise contamination rates estimated from the SpotClean model are shown in columns 1 and 2. Validation of these rates is provided in results from simulations studies shown in Supplementary Figure 12.

| Dataset | No decontamination | SpotClean | SoupX | DecontX |
|---|---|---|---|---|
| LIBD_151507 | 31.334 | **15.041** | NA | 83.016 |
| LIBD_151508 | 26.477 | **12.987** | NA | 53.596 |
| LIBD_151669 | 21.931 | **11.745** | NA | 267.124 |
| LIBD_151670 | 17.903 | **10.304** | NA | 70.851 |
| LIBD_151673 | 22.121 | **11.682** | NA | 56.747 |
| LIBD_151674 | 25.861 | **13.361** | 108.916 | 57.304 |
| mouse_brain | 24.979 | **9.896** | 779.811 | 278.374 |
| mouse_kidney | 12.114 | **7.810** | 291.890 | 119.825 |
| human_breast | 14.278 | **9.605** | 139.924 | 72.129 |
| human_lymphnode | 113.216 | **30.261** | 486.128 | 189.395 |
| human_spinalcord | 126.197 | **13.191** | 163.898 | 187.928 |

**Supplementary Table 3**: Average mean squared error (MSE) between true and decontaminated gene expression (average taken over 3000 genes) in 11 SimI datasets simulated using input from the dataset indicated. NA denotes datasets for which the corresponding method failed to run. The lowest MSE for each dataset is bolded.

| Dataset | No decontamination | SpotClean | SoupX | DecontX |
|---|---|---|---|---|
| LIBD_151507 | 30.720 | **14.957** | NA | 58.651 |
| LIBD_151508 | 26.102 | **12.909** | NA | 122.332 |
| LIBD_151669 | 21.610 | **12.001** | NA | 266.682 |
| LIBD_151670 | 17.452 | **10.221** | NA | 154.391 |
| LIBD_151673 | 21.472 | **12.172** | NA | 74.319 |
| LIBD_151674 | 25.131 | **13.469** | NA | 57.744 |
| mouse_brain | 24.161 | **9.625** | 824.134 | 284.147 |
| mouse_kidney | 12.165 | **7.903** | 319.903 | 121.810 |
| human_breast | 13.790 | **9.987** | 118.043 | 71.458 |
| human_lymphnode | 108.288 | **31.581** | 464.735 | 196.503 |
| human_spinalcord | 122.037 | **14.431** | 181.217 | 515.027 |

**Supplementary Table 4**: Average mean squared error (MSE) between true and decontaminated gene expression (average taken over 3000 genes) in 11 SimII datasets simulated using input from the dataset indicated. NA denotes datasets for which the corresponding method failed to run. The lowest MSE for each dataset is bolded.

| Dataset | No decontamination | SpotClean | SoupX | DecontX |
|---|---|---|---|---|
| LIBD_151507 | 32.472 | **14.998** | NA | 87.570 |
| LIBD_151508 | 27.371 | **13.248** | NA | 196.679 |
| LIBD_151669 | 22.892 | **11.710** | NA | 81.707 |
| LIBD_151670 | 18.538 | **10.255** | 62.989 | 63.855 |
| LIBD_151673 | 23.502 | **11.719** | NA | 184.707 |
| LIBD_151674 | 27.832 | **13.372** | NA | 61.567 |
| mouse_brain | 26.856 | **9.508** | 685.702 | 284.959 |
| mouse_kidney | 12.989 | **7.912** | 302.584 | 119.542 |
| human_breast | 15.222 | **9.953** | 135.952 | 74.705 |
| human_lymphnode | 120.495 | **28.026** | 534.534 | 195.524 |
| human_spinalcord | 133.396 | **13.414** | 186.904 | 563.552 |

**Supplementary Table 5**: Average mean squared error (MSE) between true and decontaminated gene expression (average taken over 3000 genes) in 11 SimIII datasets simulated using input from the dataset indicated. NA denotes datasets for which the corresponding method failed to run. The lowest MSE for each dataset is bolded.

| Dataset | No decontamination | SpotClean | SoupX | DecontX |
|---|---|---|---|---|
| LIBD_151507 | 36.056 | **15.388** | NA | 144.595 |
| LIBD_151508 | 30.580 | **13.639** | NA | 141.042 |
| LIBD_151669 | 25.767 | **12.677** | NA | 79.926 |
| LIBD_151670 | 21.365 | **10.553** | NA | 196.039 |
| LIBD_151673 | 27.271 | **12.769** | 97.401 | 339.906 |
| LIBD_151674 | 32.751 | **13.967** | NA | 67.959 |
| mouse_brain | 31.022 | **9.419** | 829.479 | 458.233 |
| mouse_kidney | 14.756 | **7.771** | 331.323 | 131.421 |
| human_breast | 16.731 | **9.348** | 136.516 | 76.562 |
| human_lymphnode | 132.168 | **29.893** | 523.708 | 209.594 |
| human_spinalcord | 152.092 | **13.154** | 223.067 | 215.555 |

**Supplementary Table 6**: Average mean squared error (MSE) between true and decontaminated gene expression (average taken over 3000 genes) in 11 SimIV datasets simulated using input from the dataset indicated. NA denotes datasets for which the corresponding method failed to run. The lowest MSE for each dataset is bolded.

| Dataset | Link |
| --- | --- |
| Six datasets from SpatialLIBD | https://github.com/LieberInstitute/spatialLIBD |
| mouse_brain | https://support.10xgenomics.com/spatial-gene-expression/datasets/1.1.0/V1_Adult_Mouse_Brain |
| mouse_kidney | https://support.10xgenomics.com/spatial-gene-expression/datasets/1.1.0/V1_Mouse_Kidney |
| human_breast | https://support.10xgenomics.com/spatial-gene-expression/datasets/1.1.0/V1_Breast_Cancer_Block_A_Section_2 |
| human_breast_2 | https://support.10xgenomics.com/spatial-gene-expression/datasets/1.3.0/Visium_FFPE_Human_Breast_Cancer |
| human_lymphnode | https://support.10xgenomics.com/spatial-gene-expression/datasets/1.1.0/V1_Human_Lymph_Node |
| human_spinalcord | https://support.10xgenomics.com/spatial-gene-expression/datasets/1.2.0/Targeted_Visium_Human_SpinalCord_Neuroscience |
| Two datasets from Slide-seqV2 | https://singlecell.broadinstitute.org/single_cell/study/SCP815/highly-sensitive-spatial-transcriptomics-at-near-cellular-resolution-with-slide-seqv2#study-download |

**Supplementary Table 7**: Links to 14 publicly available spatial transcriptomics datasets (12 from the 10x Visium and 2 from the Slide-seqV2 protocol).

| Gene | -log10_pval | logfc | t | CI | df |
|---|---|---|---|---|---|
| ENC1 | 21.6772 | -0.59211 | -10.0715 | -0.77218,-0.52024 | 716.6266 |
| TUBB2A | 26.82923 | -0.5005 | -11.4299 | -0.87942,-0.62153 | 611.6922 |
| MBP | 68.23467 | 0.43074 | 20.62996 | 1.31259,1.58892 | 502.7044 |
| MAP1B | 24.09136 | -0.4268 | -10.7987 | -0.90317,-0.62517 | 558.5923 |
| SNAP25 | 38.24431 | -0.42293 | -14.1973 | -1.12206,-0.84929 | 526.9102 |
| VSNL1 | 23.18776 | -0.41835 | -10.5489 | -0.85222,-0.58468 | 578.5369 |
| CCK | 30.67843 | -0.41324 | -12.3489 | -0.97287,-0.70589 | 605.6358 |
| RTN1 | 21.56528 | -0.41252 | -10.0983 | -0.77982,-0.5259 | 616.3649 |
| YWHAH | 22.66223 | -0.40974 | -10.4066 | -0.85579,-0.58405 | 586.1957 |
| OLFM1 | 21.10679 | -0.4048 | -9.99412 | -0.8003,-0.53742 | 590.4734 |

**Supplementary Table 8**: Summary statistics of t-test results for known DE genes in Figure 2b in LIBD_151507 raw data. -log10_pval: $-\log_{10}$ transformed p-value. logfc: log transformed fold change. t: t statistic. CI: 95% confidence interval. df: degrees of freedom.

| Gene | -log10_pval | logfc | t | CI | df |
|---|---|---|---|---|---|
| ENC1 | 26.81212 | -0.81797 | -11.2768 | -0.84106,-0.59168 | 828.6978 |
| TUBB2A | 36.07802 | -0.73195 | -13.3537 | -1.07278,-0.79781 | 785.5271 |
| MBP | 47.97652 | 0.458097 | 16.37517 | 1.41258,1.79774 | 507.8979 |
| MAP1B | 30.97989 | -0.64203 | -12.3232 | -1.15413,-0.83691 | 696.4673 |
| SNAP25 | 64.89745 | -0.85097 | -19.0397 | -1.70572,-1.38683 | 710.7584 |
| VSNL1 | 34.64576 | -0.70785 | -13.0801 | -1.12332,-0.83014 | 751.8196 |
| CCK | 53.54393 | -0.84157 | -16.7963 | -1.49164,-1.17946 | 779.5883 |
| RTN1 | 34.02306 | -0.69379 | -12.915 | -1.0405,-0.76594 | 787.7664 |
| YWHAH | 32.53661 | -0.6964 | -12.6217 | -1.1375,-0.83128 | 752.0688 |
| OLFM1 | 34.35643 | -0.71864 | -13.0144 | -1.08758,-0.80248 | 755.607 |

**Supplementary Table 9**: Summary statistics of t-test results for known DE genes in Figure 2b in LIBD_151507 data decontaminated by SpotClean. -log10_pval: $-\log_{10}$ transformed p-value. logfc: log transformed fold change. t: t statistic. CI: 95% confidence interval. df: degrees of freedom.

### Supplementary Section S1: Sources of contamination in RNA-sequencing experiments

We define UMI counts for a given gene in a given sample as contaminated if at least one of the counts did not arise from that gene in that sample. Common sources of contamination include barcode swapping (also called index hopping) and contamination by ambient RNA. Barcode swapping refers to a barcode from one sample binding reads from another once samples are pooled together for sequencing. Specifically, most RNA-sequencing protocols attach barcodes (sample-specific strings of nucleotides) to mRNA transcripts in a sample prior to pooling samples for sequencing so that, following sequencing, sample-specific transcript abundance can be estimated. This is the case for the 10x Visium spatial transcriptomics protocol, where mRNA binds barcodes that are spatially addressed to a given tissue spot, then are released and pooled for sequencing. The process is not without error, and numerous studies have found evidence of barcode swapping, where a barcode specific to one sample binds reads from another at random[7–10]. This is distinct from the spot swapping artifact detailed here since spot swapping is not at random. Rather, with spot swapping, the probability of a spot-specific barcode binding reads from another spot increases as the distance between spots decreases. As the statistical methods developed to adjust for barcode swapping[10] do not accommodate the spatial dependence inherent in spot swapping, they are not sufficient in this setting.

A second type of contamination is specific to scRNA-seq experiments. In droplet based scRNA-seq, for example, each droplet ideally contains one cell, and barcodes specific to that droplet bind mRNA from the cell. In practice, however, ambient (cell free) RNA may also bind barcodes from a droplet. As with barcode swapping, robust statistical methods are in place to adjust[5,6], but they are not appropriate for spatial data as they do not accommodate the spatial dependence. Supplementary Figure 13 shows results from two popular decontamination methods, SoupX[5] and DecontX[6]. Each of these methods begins by clustering single-cell data[6], or taking as input clustering information[5]; decontamination is then performed within cluster. For some marker genes that distinguish between clusters, these methods work well, or at least do not change the data very much. However, in many cases, decontamination reduces signal substantially (Supplementary Figure 13, panels a and b). In addition, since both methods decontaminate all genes within a cluster simultaneously, artificial patterns are imposed upon genes showing no spatial changes (Supplementary Figure 13, panel c). This leads to poor estimates of expression

(Supplementary Tables 3-6) and increased false discoveries in downstream analyses (Supplementary Figure 10). These results should not be taken as evidence that SoupX and DecontX perform poorly in general. That is not the case. Rather, it should be stressed neither method was designed for spatial data and, consequently, it should not be surprising that they are not sufficient in this setting.

**Supplementary Section S2: Derivation of gradients for estimating contamination parameters.**

Recall that we would like to minimize

$$\left(\widehat{r_\beta}, \widehat{r_\gamma}, \hat{\sigma}, \{\widehat{\mu_{\cdot t}}\}_{t\in I_t}\right) = \underset{r_\beta, r_\gamma, \sigma, \{\mu_{\cdot t}\}_{t\in I_t}}{\operatorname{argmin}} \sum_{j\in I_t \cup I_b} \left(X_{\cdot j} - \eta_{\cdot j}\right)^2$$

where

$$\eta_{\cdot j} = E\left(X_{\cdot j}\right) = \begin{cases} \sum_{t\in I_t} \mu_{\cdot t}\, r_\beta \left[r_\gamma \frac{1}{K} + (1 - r_\gamma) w_{t,j}\right] & \text{, if } j \in I_b \\ \mu_{\cdot j}(1 - r_\beta) + \sum_{t\in I_t} \mu_{\cdot t}\, r_\beta \left[r_\gamma \frac{1}{K} + (1 - r_\gamma) w_{t,j}\right] & \text{, if } j \in I_t \end{cases}$$

Rewriting the problem in matrix representation, let $\mu = \left(\mu_{\cdot 1}, \dots, \mu_{\cdot K_t}\right)^T$ and $X = \left(\{X_{\cdot j}\}_{j\in I_b}, \{X_{\cdot j}\}_{j\in I_t}\right)^T$. Denote the proximal contamination weight matrix $W = \begin{pmatrix} W_1 \\ W_2 \end{pmatrix}$, where $W_1$ is a $K_b \times K_t$ matrix containing the Gaussian weights between background and tissue spots, and $W_2$ is a $K_t \times K_t$ matrix containing the Gaussian weights between pairs of tissue spots. The column sums of $W$ are equal to 1 based on its definition. Let $I_{K_t}$ be the identity matrix with dimension $K_t$. Let $J_1$ and $J_2$ be $K_b \times K_t$ and $K_t \times K_t$ matrices of ones representing the distal contamination weights. The residual sum of squares (RSS) becomes

$$RSS = \left\| X - \left[(1 - r_\beta)\begin{pmatrix} 0 \\ I_{K_t} \end{pmatrix} + r_\beta(1 - r_\gamma)\begin{pmatrix} W_1 \\ W_2 \end{pmatrix} + \frac{r_\beta r_\gamma}{K}\begin{pmatrix} J_1 \\ J_2 \end{pmatrix}\right]\mu \right\|_{L_2}^2$$

The gradients of unknown parameters are then calculated as

$$\frac{\partial \text{RSS}}{\partial \mu} = 2\left[(1 - r_\beta)^2 I_{K_t} + {r_\beta}^2 (1 - r_\gamma)^2 W^T W + r_\beta(1 - r_\beta)(1 - r_\gamma)(W_2^T + W_2)\right.$$

$$\left. + \frac{r_\beta r_\gamma (2 - r_\beta r_\gamma)}{K} J_2\right]\mu$$

$$- 2\left[(1 - r_\beta)(0 \quad I_{K_t})X + r_\beta(1 - r_\gamma) W^T X + \frac{r_\beta r_\gamma}{K}\begin{pmatrix} J_1 \\ J_2 \end{pmatrix}^T X\right]$$

$$\frac{\partial RSS}{\partial r_\beta} = \mu^T \left[2(r_\beta - 1) I_{K_t} + 2 r_\beta (1 - r_\gamma)^2 W^T W + (1 - 2r_\beta)(1 - r_\gamma)(W_2^T + W_2)\right.$$

$$\left. + \frac{2r_\gamma - 2 r_\beta {r_\gamma}^2}{K} J_2\right]\mu - 2X^T\left[(1 - r_\gamma)W + \frac{r_\gamma}{K}\begin{pmatrix} J_1 \\ J_2 \end{pmatrix} - \begin{pmatrix} 0 \\ I_{K_t} \end{pmatrix}\right]\mu$$

$$\frac{\partial RSS}{\partial r_\gamma} = \mu^T \left[ 2{r_\beta}^2 (r_\gamma - 1) W^T W + r_\beta (r_\beta - 1)(W_2^T + W_2) + \frac{2r_\beta - 2{r_\beta}^2 r_\gamma}{K} J_2 \right] \mu$$

$$- 2X^T \left[ \frac{r_\beta}{K} \binom{J_1}{J_2} - r_\beta W \right] \mu$$

**Supplementary Section S3: Derivation of the EM algorithm for estimating true gene expression values.**

Recall that the observed data $\mathcal{D} = \{X_{g,j}\}_{g\in G, j\in I_t \cup I_b}$ has log-likelihood

$$l_{\mathcal{D}} = \sum_{g\in G} \sum_{j\in I_t\cup I_b} l_{X_{g,j}} = \sum_{g\in G} \sum_{j\in I_t\cup I_b} \{X_{g,j} \log \eta_{g,j} - \eta_{g,j}\} + \text{constant}$$

and the complete data are $\mathcal{C} = \{S_{g,t}, B_{g,t,j}\}_{g\in G, t\in I_t, j\in I_t\cup I_b}$ with log-likelihood

$$l_{\mathcal{C}} = \sum_{g\in G} \sum_{t\in I_t} l_{S_{g,t}} + \sum_{g\in G} \sum_{t\in I_t} \sum_{j\in I_t\cup I_b} l_{B_{g,t,j}}$$

$$= \sum_{g\in G} \sum_{t\in I_t} \left[S_{g,t} \log\left(\mu_{g,t}(1-r_\beta)\right) - \mu_{g,t}(1-r_\beta)\right]$$

$$+ \sum_{g\in G} \sum_{t\in I_t} \sum_{j\in I_t\cup I_b} \left[B_{g,t,j} \log\left(\mu_{g,t} r_\beta \left[(1-r_\gamma) w_{t,j} + r_\gamma \frac{1}{K}\right]\right)\right.$$

$$\left. - \mu_{g,t} r_\beta \left[(1-r_\gamma) w_{t,j} + r_\gamma \frac{1}{K}\right]\right] + \text{constant}$$

$$= \sum_{g\in G} \sum_{t\in I_t} \left( S_{g,t} \log\left(\mu_{g,t}(1-r_\beta)\right) + \sum_{j\in I_t\cup I_b} \left[B_{g,t,j} \log\left(\mu_{g,t} r_\beta \left[(1-r_\gamma) w_{t,j} + r_\gamma \frac{1}{K}\right]\right)\right] - \mu_{g,t} \right)$$

$$+ \text{constant}$$

Let $\{\mu_{g,t}^{(n)}\}_{g\in G, t\in I_t}$ be the parameter values at the $n$-th iteration. The E-step involves computation of the expectation of latent variables conditioning on observed data and parameter values at the current iteration. Given the fact that if $U \sim \text{Poisson}(a), V \sim \text{Poisson}(b),$ and $U$ and $V$ are independent, then $U|(U+V) \sim \text{Binomial}(U+V, \frac{a}{a+b})$, and we have

$$S_{g,t}^{(n)} \coloneqq E\left[S_{g,t} \middle| \mathcal{D}\right] = E\left[S_{g,t} \middle| X_{g,t}\right] = X_{g,t} \frac{\mu_{g,t}^{(n)}(1-r_\beta)}{\eta_{g,t}^{(n)}}$$

$$B_{g,t,j}^{(n)} \coloneqq E\left[B_{g,t,j} \middle| \mathcal{D}\right] = E\left[B_{g,t,j} \middle| X_{g,j}\right] = X_{g,j} \frac{\mu_{g,t}^{(n)} r_\beta \left[(1-r_\gamma) w_{t,j} + r_\gamma \frac{1}{K}\right]}{\eta_{g,j}^{(n)}}$$

The M-step involves maximizing the complete log-likelihood after plugging in the conditional expectations in the E-step:

$$l_{\mathcal{C}}^{(n)} = \sum_{g \in G} \sum_{t \in I_t} \left( S_{g,t}^{(n)} \log\left(\mu_{g,t}(1 - r_\beta)\right) + \sum_{j \in I_t \cup I_b} B_{g,t,j}^{(n)} \log\left(\mu_{g,t} r_\beta \left[(1 - r_\gamma) w_{t,j} + r_\gamma \frac{1}{K}\right]\right) - \mu_{g,t} \right)$$

$$\frac{\partial l_{\mathcal{C}}^{(n)}}{\partial \mu_{g,t}} = \frac{S_{g,t}^{(n)} + \sum_{j \in I_t \cup I_b} B_{g,t,j}^{(n)}}{\mu_{g,t}} - 1$$

The M-step becomes

$$\mu_{g,t}^{(n+1)} = S_{g,t}^{(n)} + \sum_{j \in I_t \cup I_b} B_{g,t,j}^{(n)}$$
